# Steady-state visually evoked potentials and feature-based attention: Pre-registered null results and a focused review of methodological considerations

**DOI:** 10.1101/2020.08.31.275602

**Authors:** Kirsten C.S. Adam, Lillian Chang, Nicole Rangan, John T. Serences

**Affiliations:** Department of Psychology, *University of California San Diego*; Institute for Neural Computation, *University of California San Diego*; Department of Neuroscience, *Georgetown University Medical Center*; Neurosciences Graduate Program, *University of California San Diego*

**Keywords:** attention, EEG, SSVEP, feature-based attention

## Abstract

Feature-based attention is the ability to selectively attend to a particular feature (e.g., attend to red but not green items while looking for the ketchup bottle in your refrigerator), and steady-state visually evoked potentials (SSVEPs) measured from the human electroencephalogram (EEG) signal have been used to track the neural deployment of feature-based attention. Although many published studies suggest that we can use trial-by-trial cues to enhance relevant feature information (i.e., greater SSVEP response to the cued color), there is ongoing debate about whether participants may likewise use trial-by-trial cues to voluntarily ignore a particular feature. Here, we report the results of a pre-registered study in which participants either were cued to attend or to ignore a color. Counter to prior work, we found no attention-related modulation of the SSVEP response in either cue condition. However, positive control analyses revealed that participants paid some degree of attention to the cued color (i.e., we observed a greater P300 component to targets in the attended versus the unattended color). In light of these unexpected null results, we conducted a focused review of methodological considerations for studies of feature-based attention using SSVEPs. In the review, we quantify potentially important stimulus parameters that have been used in the past (e.g., stimulation frequency; trial counts) and we discuss the potential importance of these and other task factors (e.g., feature-based priming) for SSVEP studies.

## Introduction

Attending to a specific feature leads to systematic changes in the firing rates of neurons that encode the relevant feature space. For example, when looking for a ripe tomato, the firing rate of neurons tuned to red will be enhanced and the firing rate of neurons tuned to other features will be suppressed (e.g. responses to green; Bartsch et al., 2017; Ipata et al., 2006; Kiyonaga & Egner, 2016; Martinez-Trujillo & Treue, 2004; Störmer & Alvarez, 2014; Y. Wang et al., 2015). Although there is broad agreement that participants may learn to suppress irrelevant distractors with sufficient experience, there is disagreement about whether these behavioral suppression effects may be volitionally implemented on a trial-by-trial basis in response to an abstract cue (i.e., a “volitional account”), or if they instead are solely implemented via implicit or statistical learning mechanisms (i.e., a “priming-based” account). Consistent with a volitional or proactive account, some work has found that participants can learn to use a trial-by-trial cue to ignore a particular color (Arita et al., 2012; Carlisle & Nitka, 2019; Chang & Egeth, 2019; Conci et al., 2019; Moher & Egeth, 2012; Reeder et al., 2017; Z. Zhang et al., 2020, for nuanced reviews, see: Geng, 2014; Van Moorselaar & Slagter, 2020). However, other work has found that a specific color needs to be repeated over many trials to be suppressed, consistent with a priming or passive account (Cunningham & Egeth, 2016; Failing et al., 2019; Geng et al., 2019; Lamy et al., 2008; Stilwell & Vecera, 2019; Theeuwes, 2013; Vatterott & Vecera, 2012; B.-Y. Won & Geng, 2020).

Although many studies have examined the effects of feature-based suppression on later selection-related event-related potential (ERP) markers such as the N2pc and Pd (Arita et al., 2012; Carlisle & Nitka, 2019; Donohue et al., 2018; Sawaki & Luck, 2010), a key open question is whether trial-by-trial cues to ignore a color modulate earlier stages of visual processing. Recent work by Reeder and colleagues (Reeder et al., 2017) hypothesized that cues about which feature to ignore (i.e., “negative cues”) may down-regulate processing in visual cortex during the pre-stimulus period. Consistent with this hypothesis, they found that overall BOLD activity in early visual cortex was lower when participants were given a negative cue about the target color than when participants were given either a positive or neutral cue. One limitation of this study, however, is that the authors were unable to test whether these univariate effects were actually feature-specific (as opposed to task-general anticipation). However, evidence from ERP studies suggests that feature-based attention modulates early visual processing in a feature-specific manner, as indexed by the P1 component (Moher et al., 2014; W. Zhang & Luck, 2009). Critically, however, these ERP studies used a design where the same target and distractor colors were repeated over many trial events. Thus, it is not clear whether feature-based attention can modulate early visual processing on a trial-by-trial basis, or if modulation of early visual processing is achieved primarily via inter-trial priming (Lamy & Kristjansson, 2013; Theeuwes, 2013). To attempt to address this gap, we conducted a pre-registered experiment in which we measured steady-state visually evoked potentials (SSVEPs) while giving participants trial-by-trial cues to suppress feature information.

When visual input flickers continuously at a given frequency (e.g., one stimulus at 24 Hz and another at 30 Hz), the visually evoked potential in the electroencephalogram (EEG) signal reflects these “steady states” and time-frequency analyses can be used to derive estimates of the strength of neural responses to each stimulus (Adrian & Matthews, 1934; Regan, 1977). The amplitude of the frequency-specific SSVEP response has been shown to be modulated by both spatial and feature-based attention (higher amplitude when attended; Chen et al., 2003; Morgan et al., 1996; Müller et al., 1998, 2006; Pei et al., 2002). When participants are cued on a trial-by-trial basis to attend to a particular feature (e.g., color), the SSVEP amplitude is higher for the attended feature (Andersen et al., 2008; Chen et al., 2003; Müller et al., 2006). Further, the time-course of the SSVEP response to an attended color reveals an early enhancement followed by a suppressed response to the irrelevant, non-attended color (Andersen & Müller, 2010; Forschack et al., 2017).

Our primary manipulation was whether we cued participants to actively attend or to actively ignore a color on a trial-by-trial basis. We planned to use this method to track enhancement vs. suppression of the SSVEP response, and to test whether the time-course of enhancement and suppression varies with cue type. In the “attend cue” condition, participants were cued about the relevant color to attend. This condition was expected to replicate prior work examining the time-course of feature-based attention using SSVEPs (Andersen & Müller, 2010), whereby enhancement of the attended color is followed by suppression of the ignored color. In the “ignore cue” condition, we instead cued participants about which color to *ignore*. If participants can use a cue to *directly* suppress a color on a trial-by-trial basis independent of target enhancement (i.e., a strong version of a volitional suppression account), we predicted that the time-course of enhancement vs. suppression of the SSVEP signal would be reduced or reversed (i.e., that suppression of the cued, to-be-ignored color may happen even *prior to* enhancement of the other color). Alternatively, if participants recode the “ignore” cue to serve as an indirect “attend” cue (e.g., “Since I’m cued to ignore blue, that means I should attend red”; Beck & Hollingworth, 2015; Becker et al., 2015; Williams et al., 2020), then we predicted that target enhancement would always precede distractor suppression regardless of whether participants were cued to attend or ignore a particular color.

To preview the results, we were unable to fully test our hypotheses about the time-course of feature-based enhancement and suppression because we did not find evidence for an overall attention effect with our task procedures. Despite robust SSVEP amplitude (Cohen’s *d* > 5), we observed no credible evidence that the SSVEP response was higher for an attended versus unattended color in either cue condition. Positive control analyses revealed that our lack of SSVEP effect was not due to a complete lack of attention to the attended color: ERP responses (P3) to the targets were modulated by attention as expected (Adamian et al., 2019; Andersen et al., 2013; Andersen, Fuchs, et al., 2011). In light of our inconclusive results, we also performed a focused methodological review of key potential task differences between our work and prior work that may have resulted in our failure to detect the effect of feature-based attention on SSVEP amplitude. We considered whether task factors such as stimulus flicker frequency, sample size, stimulus duration, and stimulus color might have impacted our ability to observe an attention effect. No single methodological factor that we considered neatly explains our lack of effect. Given our results and literature review, we propose that future work is needed to systematically explore two key factors: (1) variation in feature-based attention effects across stimulus flicker frequencies and (2) the extent to which feature-based priming modulates SSVEP attention effects.

## Methods

### Pre-registration and data availability

We published a pre-registered research plan on the Open Science Framework prior to data collection (https://osf.io/kfg9h/). Our raw data and analysis code will be made available online on the Open Science Framework at https://osf.io/ew7dv/ upon acceptance for publication.

### Participants

Healthy volunteers (*n* = 32; gender = 17 female, 15 male; mean age = 21.5 years [SD = 3.84, min = 18, max = 39]; handedness not recorded; corrected-to-normal visual acuity; normal color vision) participated in one 3.5 to 4 hour experimental session at the University of California San Diego (UCSD) campus, and were compensated $15/hr. Procedures were approved by the UCSD Institutional Review Board, and all participants provided written informed consent. Inclusion criteria included normal or corrected-to-normal visual acuity, normal color vision, age between 18 and 60 years old, and no self-reported history of major neurological disorders (e.g., epilepsy, stroke). Data were excluded from analysis if there were fewer than 400 trials in either cue condition (either due to leaving the study early or after artifact rejection). A sample size of 24 was pre-registered, and artifact rejection criteria were pre-registered (see section “EEG preprocessing” below for more details). After running each participant, we checked whether the data were usable (i.e., sufficient number of artifact-free trials) so that we would know when to stop data collection. To reach our final sample size (n = 23 participants with usable data), we ran a total of 32 participants. Nine participants’ data were not used for the following reason: Subjects with an error in the task code (n = 3), subjects who stopped the study early due to technical issues or to participants’ preferences (n = 4), subjects with too many artifacts (n = 2). Note, we were one subject short of our pre-registered target sample size of 24 because data collection was suspended due to COVID-19. However, as our later power analyses will show, we do not believe the addition of 1 further subject would have meaningfully altered our conclusions.

### Stimuli and Procedures

#### Heterochromatic flicker photometry task

We chose perceptually equiluminant colors for each participant using a heterochromatic flicker photometry task. Participants viewed a large circular, flickering stimulus (8° radius) on a black screen (0.08 cd/m2). We generated 5 circular color spaces in CIELAB-space with varying luminance (circles centered on: L = 35-65, a = 0, b = 0; 5 colors equally spaced around circle with radius = 35) for use in the task. Participants matched each of the 5 colors to a medium-gray reference color (RGB = 105.6 105.6 105.6).

On each trial, the circular background was flickered between two different colors. One color was always medium-gray, and the other color was the to-be-matched color on that trial. The colors of circular background were phase reversed at a rate of 24 Hz, giving the appearance of a fast flicker when the subjective luminance values were not matched. On top of the flickering circular stimulus small oriented bars were drawn in the medium-gray reference color (the bars changed locations at a rate of 1Hz). The oriented bars served no purpose other than subjectively making it easier to discriminate fine-grained differences in luminance between the flickering colors (i.e., these bars gave secondary visual cues about equiluminance via the “minimally distinct border” phenomenon, Kaiser, 1988). Participants increased or decreased the luminance of the to-be-matched color (using up and down arrow keys) until the amount of perceived flicker was minimized – the point of perceptual equiluminance. The luminance starting value of the to-be-matched color was chosen at random on each trial. Once satisfied with their response, the participant pressed spacebar to continue to the next trial. Each to-be-matched color was repeated 3 times (15 trials total).

#### Feature-based attention task

All stimuli were viewed on a luminance calibrated CRT monitor (1024 x 768 resolution, 120 Hz refresh rate) from a distance of ~50 cm in a dimly lit room. Stimuli were generated using Matlab 2016a and the Psychophysics toolbox (Brainard, 1997; Kleiner et al., 2007; Pelli, 1997). Participants rested their chin on a chin-rest and fixated a central dot (0.15° radius) throughout the experiment. The stimulus was a circular aperture (radius = ~9.5°) filled with 120 oriented bars (each bar ~1.1° long and ~.1° wide). Bars were centered on a grid and separated by ~1 bar length such that they never overlapped with one another. On each individual frame (~8.33 ms) this grid was randomly phase shifted (0:2π in x and y coordinates) and rotated (1:360°), thus giving the appearance of random flicker. To achieve Steady State Visually Evoked Potentials (SSVEP) half of the bars flickered at 24 Hz (3 frames on, 2 frames off) and the other half flickered at 30 Hz (2 frames on, 2 frames off). Due to the jittered rotation of bar positions and to the random assignment of colors to bars on each “on” frame, this means that the individual pixels that were “on” for each color varied from frame to frame. The unpredictable nature of each bar’s exact position is thus quite similar to unpredictable stimuli that have been used in past work (e.g., Andersen et al., 2008). For each “off frame” no bars of that color were shown (e.g., if 24 Hz had an “on” frame and 30 Hz had an “off” frame, then only 60 out of 120 bars would be shown on the black background). If both the 24 Hz and 30 Hz bars were “off”, then a black screen would be shown on that frame. See Appendix A for an illustration of some example frame-by-frame screenshots of the stimuli.

On each trial (Figure 1), the participants viewed the stimulus array of flickering, randomly oriented bars presented on a black background (0.08 cd/m2). Half of these bars were shown in one color (randomly chosen from the 5 possible colors) and the other half were in another randomly chosen color (with the constraint that the two sets of bars must be two different colors). During an initial baseline (1,333 ms), participants viewed the flickering dots while they did not yet know which color to attend; during this baseline, the fixation point was a medium gray color (same as the reference color in the flicker photometry task). After the baseline, the fixation dot changed color, cuing the participants about which color to attend or ignore. In the “attend cue” condition, the color of the fixation point indicated which color should be attended. In the “ignore cue” condition, the color of the fixation point indicated which color should be ignored. These two conditions were blocked, and the order was counterbalanced across participants (further details below). During the stimulus presentation (2,000 ms), participants monitored the relevant color for a brief “target event” (333 ms). During this brief target event, a percentage of lines in the relevant color will be coherent (iso-oriented). Critically, the orientation of each coherent target or distractor event was completely unpredictable (randomly chosen between 1-180 degrees); thus, participants could not attend to a particular orientation in advance in order to perform the task. A target event occurred on 50% of trials, and participants were instructed to press the spacebar as quickly as possible if they detected a target event. Importantly, physically identical events (iso-oriented lines in a random orientation, 333 ms) could also appear in the distractor color (50% of trials). Participants were instructed that they should only respond to target events; if they erroneously responded to the distractor event, the trial was scored as incorrect. The target and/or distractor events could begin as early as cue onset (0 ms) and no later than 1,667 ms after stimulus onset). Participants could make responses up to 1 second into the inter-trial interval. If both a target and distractor event were present, their onset times were separated by at least 333 ms.

**Figure 1.**
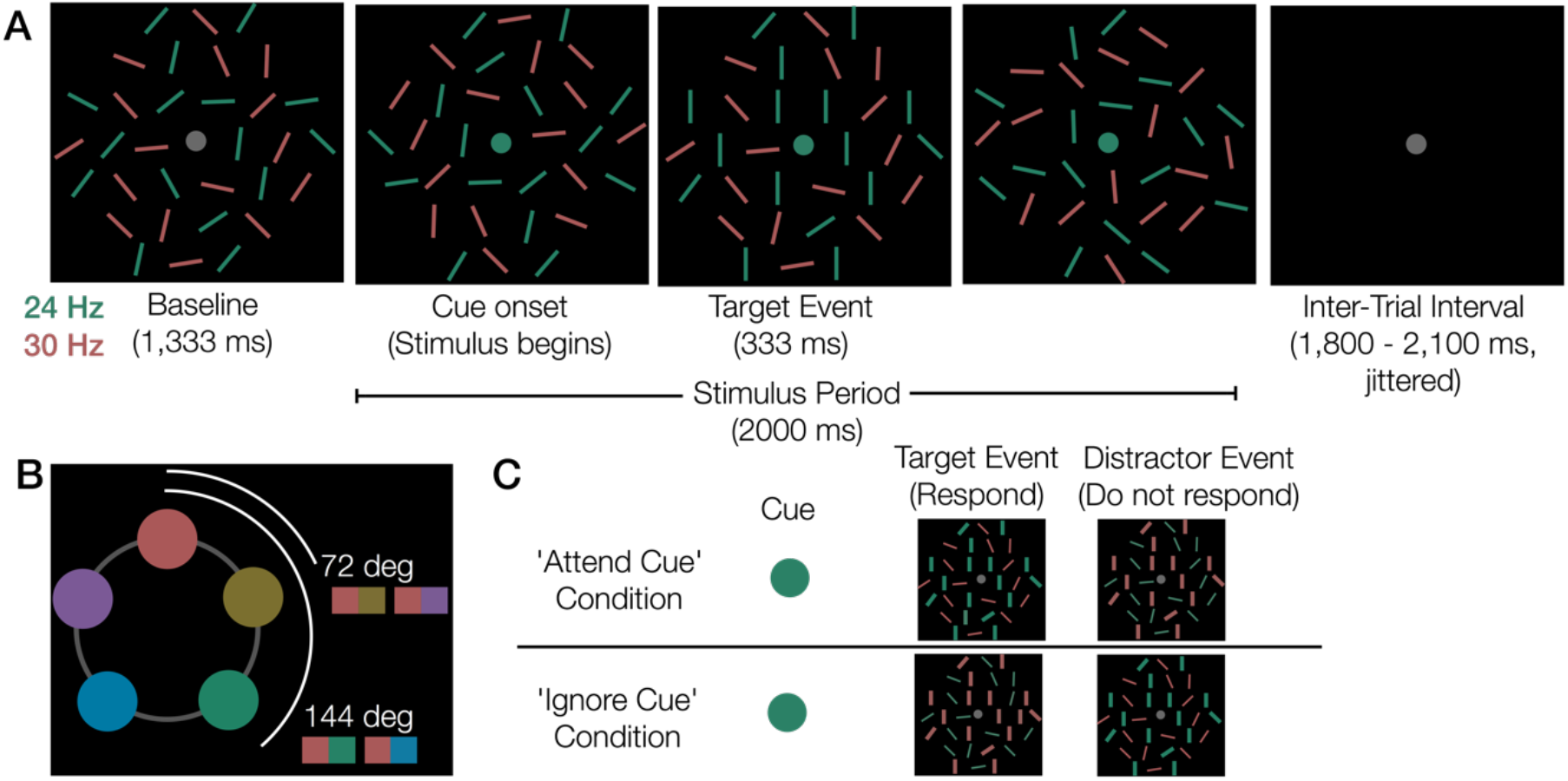
Feature-based attention task. (A) Trial events in an example ‘attend cue’ trial where there is a target event. Note, this figure is a schematic and the stimuli are not drawn to scale. After a baseline period, participants were cued to attend or ignore one color via a change to the fixation point color. If participants noticed a target event (~75% iso-oriented lines in the to-be-attended color), they pressed the space bar. (B) Five colors were used, and these 5 colors appeared with equal probability. Thus, the target and distractor colors could be either 72 degrees or 144 degrees apart on a color wheel. This figure shows all possible color pairings if red was the target color. (C) Examples of cues, target events, and distractor events in the 2 main conditions. In the ‘attend cue’ condition, participants made a response when the iso-oriented lines were the same color as the cue (target event) and did not respond if the iso-oriented lines occur on the uncued color (distractor event). In the ‘ignore cue’ condition, participants made a response when the iso-oriented lines occurred on the uncued color (target event), and they did not respond if the iso-oriented lines occurred on the cued color (distractor event). Note, all lines were of equal size in the real experiment; lines are shown at different widths here for easier visualization of the target and distractor colors. Here, the iso-oriented lines are drawn at vertical in all 4 examples. In the actual task, the iso-oriented lines could be any orientation (1-180).

To ensure that the task was effortful for participants, the coherence of the lines in the target stimulus was adapted at the end of each block if behavior was outside the range of 70 - 85% correct. At the beginning of the session, the target had 50% coherent iso-oriented lines. If accuracy over the block of 80 trials was >85%, coherency decreased by 5%. If block accuracy was <70%, coherency increased by 5%. The maximum allowed coherence was 80% iso-oriented lines (so that participants would not be able to simply individuate and attend a single position to perform the task) and the minimum allowed coherence was 5%. The presence and absence of target and distractor events was balanced within each block yielding a total of 4 sub-conditions within each cue type (25% each): (1) target event + no distractor event, T1D0 (2) no target event + distractor event, T0D1 (3) target event + distractor event, T1D1 (4) no target event + no distractor event, T0D0.

Participants completed both task conditions (attend cue and ignore cue). The two conditions were blocked and counterbalanced within a session (i.e., half of participants performed the “attend cue” task for the first half of the session and the “ignore cue” task for the second half of the session.) Each block of 80 trials took approximately 6 min 50 sec. Participants completed 18 blocks (9 per condition) for a total of 720 trials per cue condition. Note, we originally planned for 20 blocks (10 per condition) in the pre-registration, but the block number was reduced to 18 after the first few participants did not finish all blocks.

#### Summary of deviations from the registered procedures

As described in-line above, there were some minor deviations from the pre-registration: (1) We made changes to the pre-registered task code to fix errors that we discovered while running the first 3 subjects (e.g., incorrect cues and behavioral feedback in the ‘ignore cue’ condition). (2) We included code for eye-tracking, which allowed us to give participants automated real-time feedback if they blinked when they were not supposed to, i.e., during the stimulus period. (3) We reduced the total number of experimental blocks from 20 (10 per cue condition) to 18 (9 per cue condition) due to time constraints. (4) We had to prematurely stop data collection at n = 23 out of 24 due to COVID-19. (5) We forgot to specify a specific statistical test for quantifying the robustness of overall SSVEPs in section “Checking that an SSVEP is elicited at the expected frequencies before collecting the full sample”, so we have described our justification for the statistical tests we present here. (6) Due to unanticipated failure to detect an overall attention effect, we performed additional non-pre-registered control analyses to attempt to rule out possible explanations of this null effect (see section: “Non pre-registered control analyses” below).

### EEG pre-processing

Continuous EEG data were collected online from 64 Ag/AgCl active electrodes mounted in an elastic cap using a BioSemi ActiveTwo amplifier (Cortech Solutions, Wilmington, NC). An additional 8 external electrodes were placed on the left and right mastoids, above and below each eye (vertical EOG), and lateral to each eye (horizontal EOG). Continuous gaze-position data were collected from an SR Eyelink 1000+ eye-tracker (sampling rate: 1,000 Hz; SR Research, Ottawa, Ontario). We also measured stimulus timing with a photodiode affixed to the upper left-hand corner of the monitor (a white dot flickered at the to-be-attended color’s frequency; the photodiode and this corner of the screen were covered with opaque black tape to ensure it was not visible). Data were collected with a sampling rate of 1024 Hz and were not downsampled offline. Data were saved unfiltered and unreferenced (see: Kappenman & Luck, 2010), then referenced offline to the algebraic average of the left and right mastoids, low-pass filtered (<80 Hz) and high-pass filtered (>.01 Hz). Artifacts were detected using automatic criteria described below, and the data were visually inspected to confirm that the artifact rejection criteria worked as expected. We excluded subjects with fewer than 400 trials remaining per cue condition.

#### Eye movements and blinks

We used the eye-tracking data and the HEOG/VEOG traces to detect blinks and eye movements. Blinks were detected on-line during the task using the eye tracker. If a blink was detected (i.e., missing gaze position returned from the eye tracker), the trial was immediately terminated and the participant was given feedback that they had blinked (i.e., the word “blink” was written in white text in the center of the screen). If eye-tracking data could not be successfully collected (e.g., calibration issues), the VEOG trace was used to detect blinks and/or eye movements during offline artifact rejection. To do so, we used a split-half sliding window step function (Luck, 2005; window size = 150 ms, step size = 10 ms, threshold = 30 microvolts.) We also used a split-half sliding-window step function to check for eye-movements in the gaze-coordinate data from the eye-tracker (window size = 80 ms, step size = 10 ms, threshold = 1°) and in the horizontal electrooculogram (HEOG), window size = 150 ms, step size = 10 ms, threshold = 30 microvolts, and to detect blinks and/or eye movements in the vertical electrooculogram (VEOG),

#### Drift, muscle artifacts, and blocking

We checked for drift (e.g., skin potentials) by comparing the absolute change in voltage from the first quarter of the trial to the last quarter of the trial. If the change in voltage exceeded 200 microvolts, the trial was rejected for drift. In addition to slow drift, we also checked for sudden, step-like changes in voltage with a sliding window (window size = 250 ms, step size = 20 ms, threshold = 200 microvolts). We excluded trials for muscle artifacts if any electrode had peak-to-peak amplitude greater than 200 microvolts within a 15 ms time window (step size = 10 ms). We excluded trials for blocking if any electrode had ~120 ms during which all values within 1 microvolt of each other (sliding 200 ms window, step size = 50 ms).

### Pre-registered SSVEP analyses

#### General method for SSVEP quantification

We planned to quantify the SSVEP response by filtering the data with a Gaussian wavelet function. First, we calculated an average ERP for each condition at electrodes of interest (O1, Oz, O2). We chose these 3 electrodes based on large SSVEP modulations at these sites in prior work (e.g., Itthipuripat et al., 2013; Müller et al., 2006). After calculating an ERP for each condition, we filtered the data with a Gaussian wavelet functions (frequency-domain) with .1 fractional bandwidth to obtain frequency-domain coefficients from 20 to 35 Hz in 1-Hz steps. The frequency-domain Gaussian filters thus had a full-width half maximum (FWHM) that varied according to frequency, as specified by the 0.1 fractional bandwidth parameter (e.g., 1 Hz filter = FWHM of .1 Hz, 20 Hz filter = FWHM of 2 Hz, 30 Hz filter = FWHM of 3 Hz, etc). For a similar analytic approach see: (Canolty et al., 2006; Itthipuripat et al., 2013; Rungratsameetaweemana et al., 2018). Signal-to-noise ratio for each SSVEP frequency was calculated as the power at a given frequency divided by the average power of the 2 adjacent frequencies on each side. For example, SNR of 24 Hz would be calculated as the power at 24 Hz divided by the average power at 22, 23, 25, and 26 Hz. We also pre-registered an analysis plan for examining the time-course of SSVEP amplitude. However, because our data failed to satisfy pre-registered pre-requisite analyses, we do not report these time-course effects here (for completeness, we show the time-course of SNR in Figure S3).

#### Checking that an SSVEP is elicited at the expected frequencies before collecting the full sample

At n = 5, we planned to confirm that our task procedure successfully produced reliable SSVEP responses (i.e., check that we observed peaks at the correct stimulus flicker frequencies). If our task procedures failed to elicit an SSVEP at the expected frequencies, we had planned to stop data collection and alter the task to troubleshoot the problem (e.g., optimize timing, choose different flicker frequencies, make stimuli brighter, etc.). We planned to begin data collection over again if we failed this trouble-shooting step. Note, at this early stage we only verified if the basic method worked (SSVEP frequencies were robust): we did not test whether any hypothesized attention effects were present, as this could inflate our false discovery rate (Kravitz & Mitroff, 2017). Note, in the original pre-registration we failed to specify what test we would run to determine if SSVEP frequencies were robustly represented in the EEG signal. Theoretical chance for SNR would be 1, so the simplest test would be to compare the SNR for our stimulation frequencies (24 and 30 Hz) to 1 using a t-test, which we report. However, it is often is better to compare to an empirical baseline with a reasonable amount of noise (Combrisson & Jerbi, 2015). As such, we opted to also compute an effect size comparing the SNR for our stimulation frequencies to all other frequencies (with the exception that we did not use frequencies +/- 2 Hz of 24 or 30 Hz as baseline values, since SNR was calculated as the power at frequency F divided by the power in the 2 adjacent 1-hz bins).

#### Checking achieved power for the basic attention effect

Without adequate power for the basic attention effect, we would not be able to robustly interpret the time-course of enhancement vs. suppression. At the full sample size, we thus planned to check whether we had sufficient achieved power for the overall attention effect (>=80% power for attended vs. ignored color collapsed across the entire stimulus period) as a prerequisite for interpreting the time-course of enhancement and suppression.

#### Checking if a priori electrodes are reasonable

We chose a priori to analyze the SSVEP at electrodes O1, Oz, and O2 (Itthipuripat et al., 2013; Rungratsameetaweemana et al., 2018), but we planned to plot the topography of SSVEP modulation across all electrodes to check that these a priori electrodes were responsive to the SSVEP manipulation. If these electrodes were not responsive to the SSVEP, we planned to perform a cluster-based permutation test to select a new set of electrodes.

#### Checking if target and/or distractor presence alters results

Our core analyses planned to use all trials for each condition (e.g., target event present or absent, distractor event present or absent). To confirm that the act of making a response did not contaminate the SSVEP results, we planned to compare SSVEPs for each of the 4 sub-conditions (the 4 possible combinations of target present/absent and distractor present/absent). We predicted that the main SSVEP attention effect (entire stimulus period) would not be different across these 4 conditions. But, if we found an effect of target or distractor presence on the main SSVEP attention effect, then we planned to use only the trials without target or distractor events for the time-course analyses (as has been done in prior work, e.g., Andersen & Müller, 2010).

#### Checking if the distance between the target and distractor color alters results

We planned to test whether the magnitude and time-course of attentional selection differs as a function of target-distractor color similarity (~72° vs. ~144° separation between the attended and ignored color). Prior work found that it was more difficult to simultaneously attend opposite colors (180° apart) than to simultaneously attend two moderately-spaced colors (60° apart) (Chapman et al., 2019; Geweke et al., 2018; Störmer & Alvarez, 2014). Given this result, we predicted that it should be easier to suppress a color that is drastically different from the target color (and behavioral accuracy should likewise be higher for the 144° condition).

### Additional non pre-registered SSVEP and ERP control analyses

We did not anticipate our failure to find an overall attention effect with these task procedures and set of pre-registered “sanity check” analyses described above. To further understand the lack of SSVEP attention effect, we performed additional non-pre-registered control analyses.

#### Positive control: Frequency analysis of the photodiode voltage

During the recording, a photodiode was used to ensure that the flicker frequencies were faithfully presented, and the photodiode recorded voltage fluctuations induced by a small white dot that flickered at the to-be-attended target frequency on each trial. The electrical activity from the photodiode was recorded as an additional “electrode” in the data matrix (with the structure: trials x electrodes x timepoints). Thus, we performed a fast Fourier transform (FFT, Matlab function “fft.m”) to ensure that the trial indexing and FFT aspects of our analysis were correct. If these aspects of the analysis were correct, we should expect a near-perfect modulation of photodiode FFT amplitude by attention condition as only the flicker frequency of the attended stimulus was tagged. For 1 subject, the photodiode was not plugged in (leaving 22 subjects for this analysis).

#### Analysis control: Using a more similar frequency analysis procedure to prior published work

Because we were interested in characterizing a time-course effect, we chose to use a Gaussian wavelet procedure to quantify power for each frequency of interest. However, given the lack of overall attention effect, we were not able to meaningfully look at the time-course effects. Thus, we additionally used a fast Fourier transform (FFT) to measure SSVEP amplitude during the entire stimulus period. This method is more commonly used in prior published work (e.g., Andersen et al., 2008; Andersen & Müller, 2010). Following prior work, we used the entire stimulus epoch starting from 500 ms onward (500 ms – 2000 ms), we detrended the data, and we zero-padded this time window (2,048 points) to precisely estimate our frequencies.

#### Positive control: Analysis of event-related potential (P3) for an attention effect

Prior work has found that attention-related SSVEP modulations are accompanied by changes to event-related potentials (ERPs) associated with target selection and processing. To measure the P3, we calculated event-related potentials time-locked to the target or distractor onset (baselined to −200 ms to 0 ms relative to target onset). We included trials where there was only one target or distractor event on that trial, to avoid the possibility of overlap between the two signals. We calculated P3 voltage at electrodes Pz and POz during the time window 450-700 ms after target onset, similar to prior work (Adamian et al., 2019; Andersen et al., 2013; Andersen, Fuchs, et al., 2011).

## Results

### Behavior

Participants were overall accurate at the task (percent correct = 65.7%, d-prime = 1.25), and were significantly above chance (percent correct >50%, *p* < .001; d-prime >0, *p* < .001). There was no overall significant effect of cue condition (attend cue versus ignore cue) on performance, *p* = .85, but analysis of target-present trials suggested that participants could more quickly use attend cues than ignore cues (p < .05 when the target appeared between 0 ms and 275 ms, but p > .05 if the target appeared after 275 ms; Appendix B).

Average percent correct was lower than our pre-specified target range of 70-85%, meaning that most participants saw targets and distractors that were maximally coherent (80% iso-oriented lines) for the majority of blocks (mean coherence of the target/distractor events = 73.7%, SD = 5.08%). Although overall accuracy was slightly out of the range we had expected when planning the study, participants were well above chance and they saw targets with coherence values typical of prior work (Andersen et al., 2008; Andersen, Fuchs, et al., 2011; Andersen & Müller, 2010).

### Pre-registered SSVEP results

We first confirmed that our SSVEP procedure was effective at eliciting robust, frequency-specific modulations of the EEG signal. After collecting the first 5 participants, we checked that overall SSVEP amplitudes for our two target frequencies (24 and 30 Hz) were robust when collapsed across conditions (Fig 2A) before proceeding with data collection. We indeed found that the SSVEP signal was robust during the stimulus period even with n=5 for both the 24 Hz frequency (mean SNR = 4.45, SD = .14, SNR > 1: *p* < .001) and for the 30 Hz frequency (mean SNR = 2.97, SD = .15, SNR > 1: *p* < .001). These values were similar for the full n=23 sample (Fig 2B). To compute an effect size, we compared SNR values for each target frequency (24 Hz and 30 Hz) to the SNR values for each baseline frequency (frequencies from 3-33 Hz not within +/-2 Hz of 24 or 30 Hz). SNR values for the target frequencies were significantly higher than baseline, mean Cohen’s d = 5.10 (SD = 1.11) and 5.99 (SD = 2.66), respectively (See Appendix C). As planned, we also confirmed that the electrodes we selected *a priori* (O1, Oz, and O2) were reasonable given the topography of overall SSVEP amplitudes (i.e., they fell approximately centrally within the brightest portion of the heat map; Figure 2C-D).

**Figure 2.**
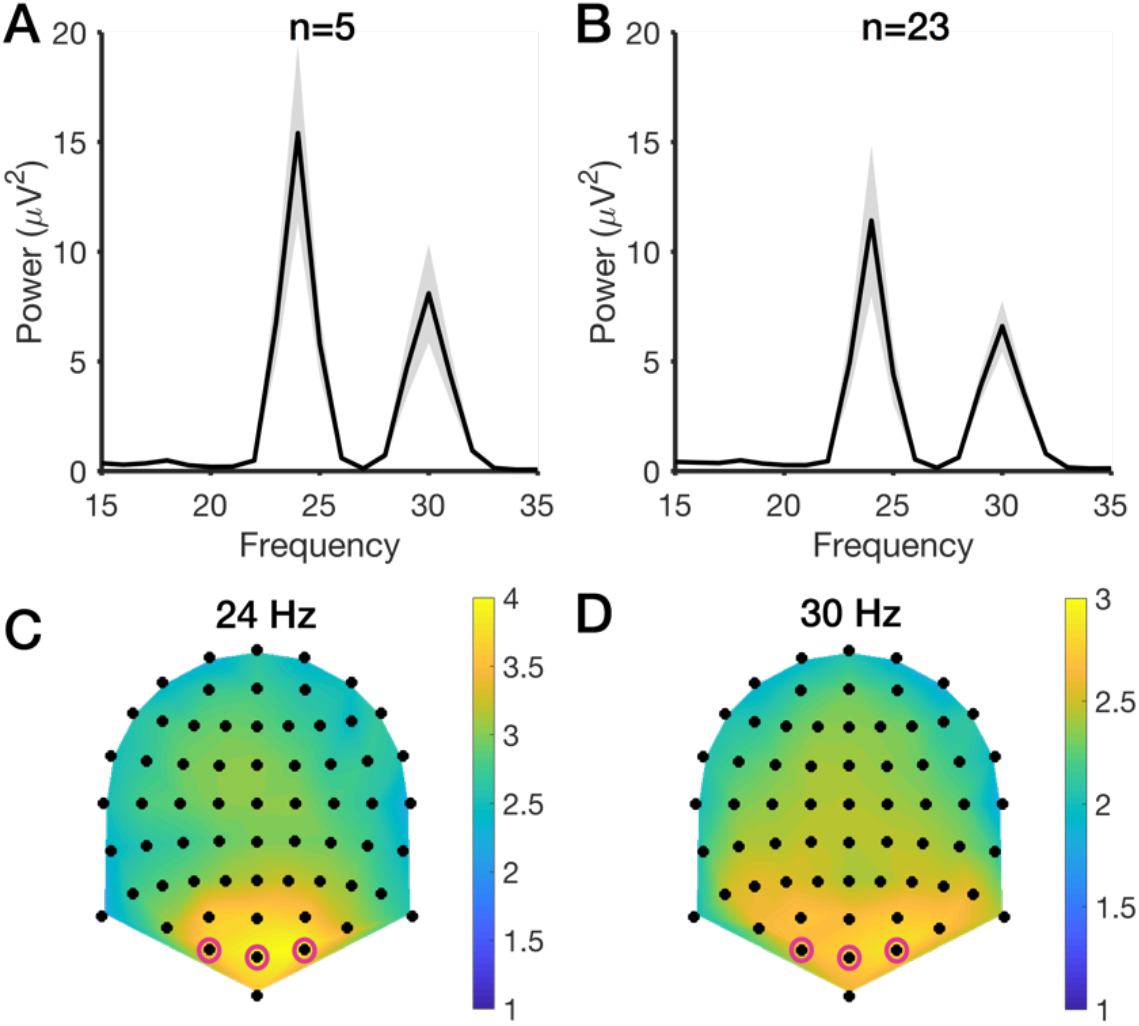
SSVEP amplitude at the expected frequencies (collapsed across all experimental conditions). (A) Power as a function of frequency during the stimulus period at electrodes O1, O2, and Oz for the first 5 participants. As expected, we observed robust peaks at the stimulated frequencies (24 and 30 Hz). (B) Power as a function of frequency for the stimulus period for the full sample (n=23). (C-D) Topography of signal to noise ratio values for 24 Hz (C) and 30 Hz (D) for all participants collapsed across all experimental conditions. Color scale indicates SNR. As expected, the *a priori* electrodes O1, O2, and Oz (magenta circles) showed robust SNR during the stimulus period.

Next, we checked for a basic attention effect, defined as a larger amplitude response evoked by the attended frequency compared to the ignored frequency). Note, for the sake of clarity, all conditions are translated into “attend” terminology. That is, if a participant was cued to “ignore blue” (24 Hz) during the “ignore cue” condition (and the other color was red and 30 Hz), this will instead be plotted as “attend red” (30 Hz). Figure 3 shows the Gaussian wavelet-derived frequency spectra during the stimulus period (500-2000 ms) as a function of cue type (attend versus ignore) and attended frequency (attend 24 Hz or attend 30 Hz). We found a main effect of measured frequency, whereby SNR was overall higher for 24 versus 30 Hz, F(1,22) = 57.89, *p* < .001, η^2^p = .73. However, we found no main effect of attended frequency (*p* = .27) or cue type (*p* = .83), and we found no significant interactions (*p* >= .18). Collapsed down to a paired t-test, the observed effect size for attended versus unattended SNR values was Cohen’s *d =* .03. To detect an effect of this size with 80% power (1-β = .8; α = .05) would require a sample size *n* > 7,000^*^. Given that we did not find an overall attention effect, we did not analyze or interpret analysis of the SSVEP time-course. However, for completeness we have shown the time course in Appendix D.

**Figure 3.**
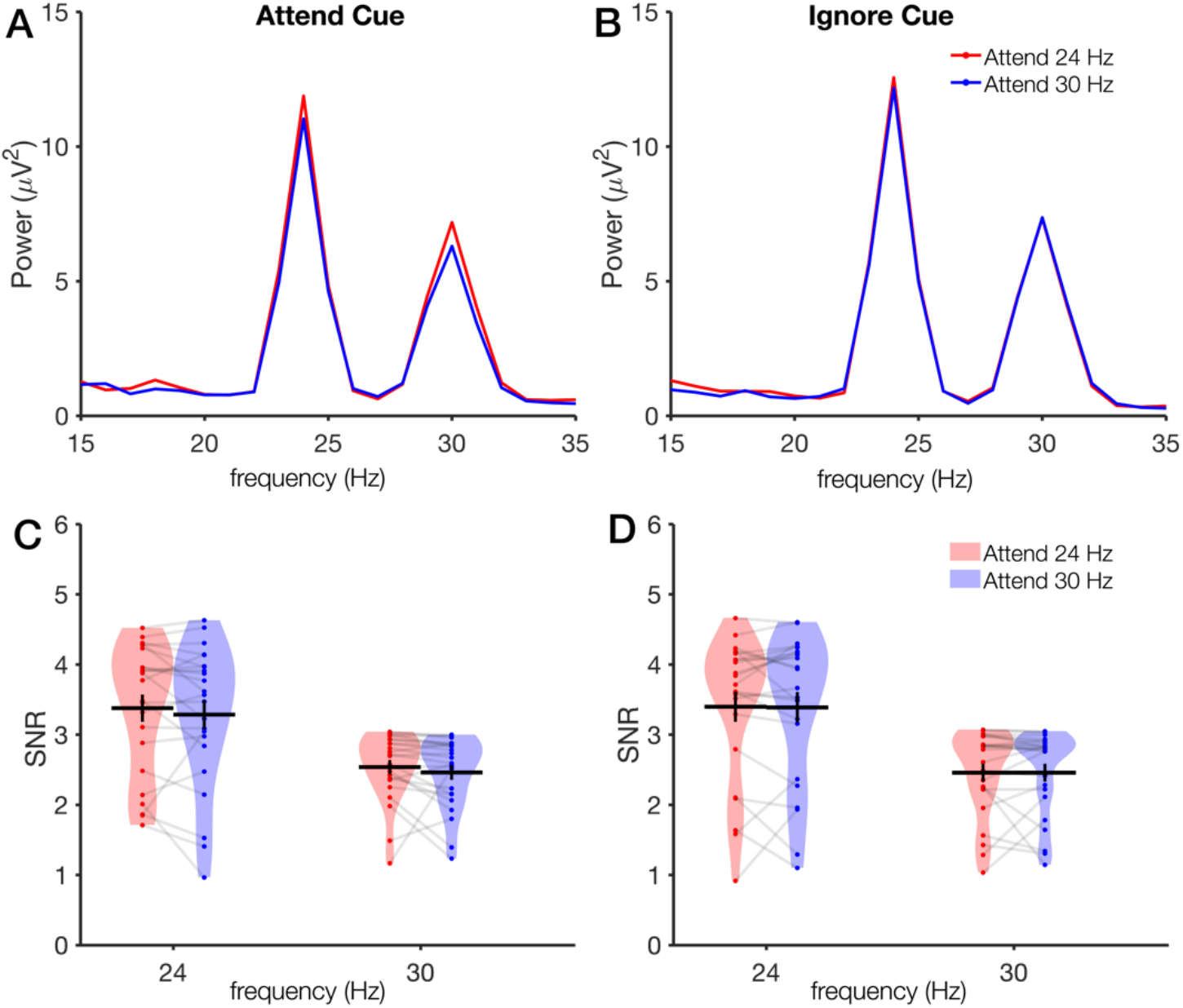
Overall attention effect in the attend cue and ignore cue conditions. (A-B) Frequency spectra in the attend cue (A) and ignore cue (B) conditions during the stimulus period. Although we observe expected peaks at 24 Hz and 30 Hz, this SSVEP response is not modulated by the attention manipulation. (C-D). Violin plots of the signal-to-noise ratio at the SSVEP frequencies in the attend cue (C) and ignore cue (D) conditions.

Although we pre-registered that we would analyze all trials (those with and without target/distractor events), most prior studies have included only trials without any target or distractor events in the main SSVEP analysis (e.g., Andersen et al., 2008; Müller et al., 2006). To ensure that our null result was not due to this analysis choice, we also planned in our pre-registration to examine the SSVEP attention effect for trials with and without target and distractor events. When restricting our analysis to only trials without targets or distractors (25% of the 1440 trials, or 360 trials total before artifact rejection), we likewise found no attention effect. As before, we found a main effect of measured frequency (24 > 30 Hz), *p* < .001, but no effect of cue condition (*p* = .053) or attended frequency (*p* = .073), and, most critically, we found no interaction between measured frequency and attended frequency (*p* = .33). Frequency spectra for all combinations of target and distractor presence are shown in Appendices E and F.

Finally, we also pre-registered that we would check whether the similarity of the target and distractor colors (72 versus 144 degrees apart on a circular color wheel; Figure 1B) would modulate the SSVEP attention effect. We likewise found that the similarity of the distractor colors did not significantly modulate the SSVEP response, and we found no attention effect (interaction of measured frequency and attended frequency) in either color distance condition (*p* >= .26; Appendix G).

### Non-pre-registered control analyses

We conducted additional control analyses to rule out possible sources of our failure to find an attention effect. First, we examined the photodiode recording to rule out any failures due to trial indexing. The photodiode measured the luminance of a white dot that flickered at the attended frequency on each trial. As expected, performing an FFT on the photodiode time-course thus yielded near-perfect tracking of the attended frequency (Figure 4A-B, *p* < .001). On the other hand, we again found null results for the main attention manipulation (Figure 4C-F) when using an FFT analysis that more closely followed prior work. We ran a repeated measures ANOVA on the signal to noise ratio values during the stimulus period, including the factors Measured Frequency (24 Hz, 30 Hz), Attended Frequency (24 Hz, 30 Hz), and Cue Type (Attend, Ignore). We found no main effect of measured frequency (*p* = .91), attended frequency (*p* = .45), or cue type (*p* = .54), and we found no significant interactions (*p* >= .38). However, the average signal-to-noise ratio of the stimulus frequencies was overall robust (M = 4.45, SD = 1.45, greater than chance value of 1: *p* < 1×10^-9^), so our inability to observe the attention effect was not due to lack of overall SSVEP signal. Likewise, we ran an additional analysis to ensure that our choice of pre-registered choice of SNR measure did not explain our null result (Appendix H).

**Figure 4.**
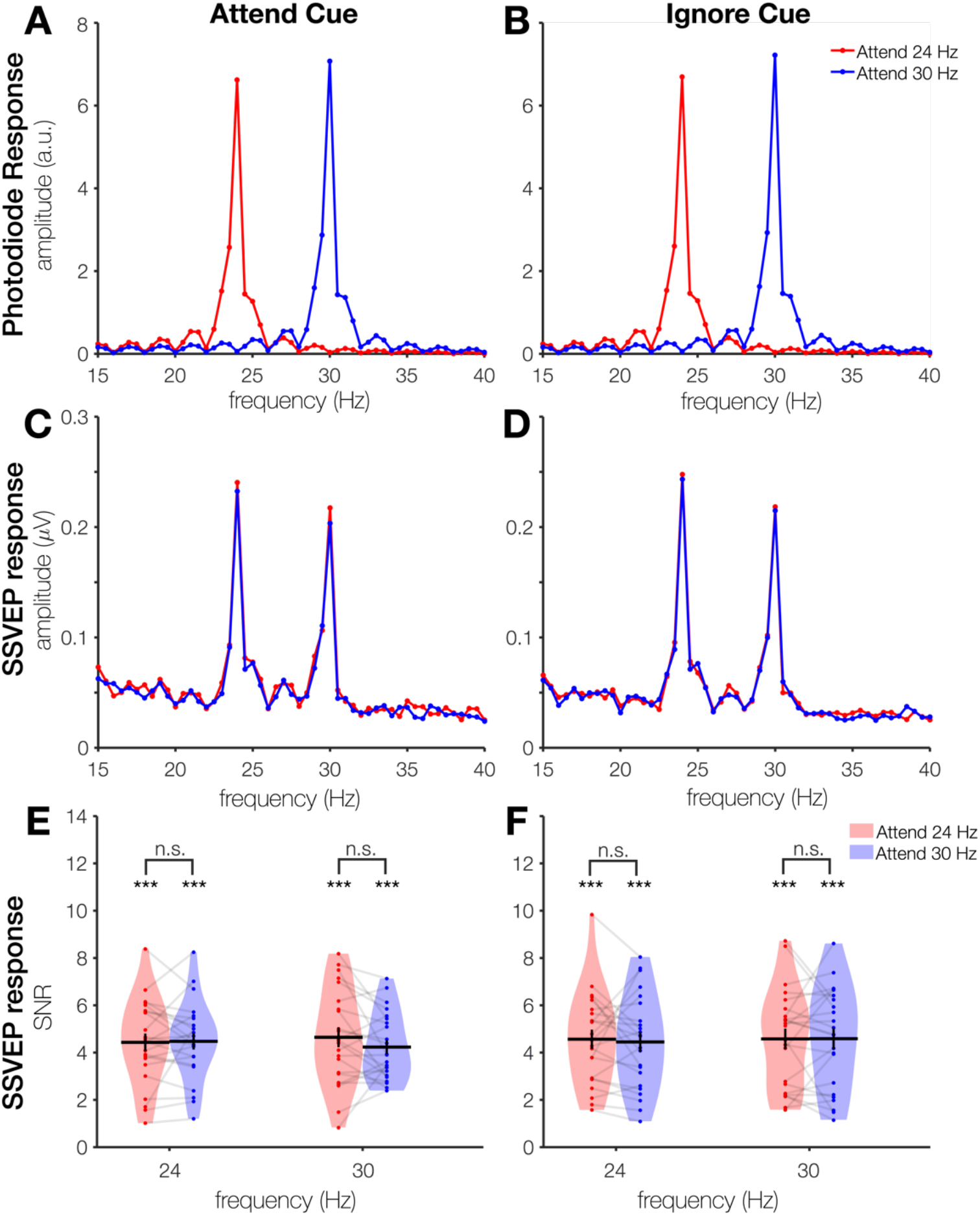
FFT analysis of the photodiode and SSVEPs at attended and unattended frequencies during the stimulus period (500-2000 ms). (A-B) As a positive control for our analysis pipeline, we performed an FFT analysis on the photodiode trace. The photodiode recorded a flickering white dot at the attended frequency on each trial. As expected, this provides a near-perfect tracking of the attended frequency in both the attend cue condition (A) and the ignore cue condition (B). (C-F) To ensure our null effect was not due to using Gaussian wavelets rather than an FFT, we repeated the main analysis with an FFT. Frequency spectra for the attend cue condition (C) and ignore cue condition (D) reveal an overall robust SSVEP signal at 24 Hz and 30 Hz, but no modulation by attention. Likewise, violin plots of signal-to-noise ratios again show robust signal but no modulation by attention in either the attend cue condition (E) or the ignore cue condition (F).

Given that some work has reported significant effects only for the second harmonic (e.g., Kim et al., 2007; Vissers et al., 2017), we likewise examined SSVEP amplitude at 48 Hz and 60 Hz, with the caveat that the 60 Hz harmonic is contaminated by line noise (Appendices I and J). We found no significant attention effects for either second harmonic frequency. We also re-ran the FFT analysis with other electrode-selection methods to ensure our *a priori* choice of electrodes did not impede our ability to observe an effect. We found no evidence that electrode choice led to our null effect, as exploiting information from all 64 electrodes by implementing rhythmic entrainment source separation (RESS) likewise yielded null effects (Appendices K and L; M. X. Cohen & Gulbinaite, 2017). To ensure that inconsistent task performance did not lead to null effects, we repeated the main FFT analysis on only accurate trials. We likewise found null attention effects when analyzing only accurate trials (Appendix M).

Finally, we tested whether phase consistency, rather than power, may track attention in our task (e.g., Nunez et al., 2015; Tallon-Baudry et al., 1996). To do so, we performed an FFT on single trials rather than on condition-averaged waveforms, and we extracted single-trial phase values. We calculated a phase-locking index by computing mean-resultant vector length on each condition’s histogram of single-trial phase values. Mean-resultant vector length ranges from 0 (fully random values) to 1 (perfectly identical values), for reference, see Zar (2010). We found no effect of attention on this phase-locking index (Appendix N).

### Positive control: Analysis of event-related potential (P3b) for an attention effect

Consistent with prior work, we found a significantly larger P3 component for target onsets compared to distractor onsets (Figure 5). A repeated measures ANOVA with within-subjects factors cue type (attend cue or ignore cue) and event type (target or distractor onset) revealed a robust main effect of event type (target > distractor), *F*(1,22) = 51.64, *p* < 1×10^-5^, η^2^_p_ = .70, and a main event of cue type (attend > ignore), F(1,22) = 4.96, *p* = .037, η^2^_p_ = .18, but no interaction between event type and cue type (*p* = .65). Control analyses confirmed this P3 modulation was not due to differential rates of making a motor response for targets and distractors (Appendix O). The main effect of event type (target > distractor) remained when analyzing only trials where participants made a motor response (*p* < .001). Thus, the P3 was overall larger for target than distractor events, consistent with prior work that found this ERP attention effect alongside an SSVEP attention effect.

**Figure 5.**
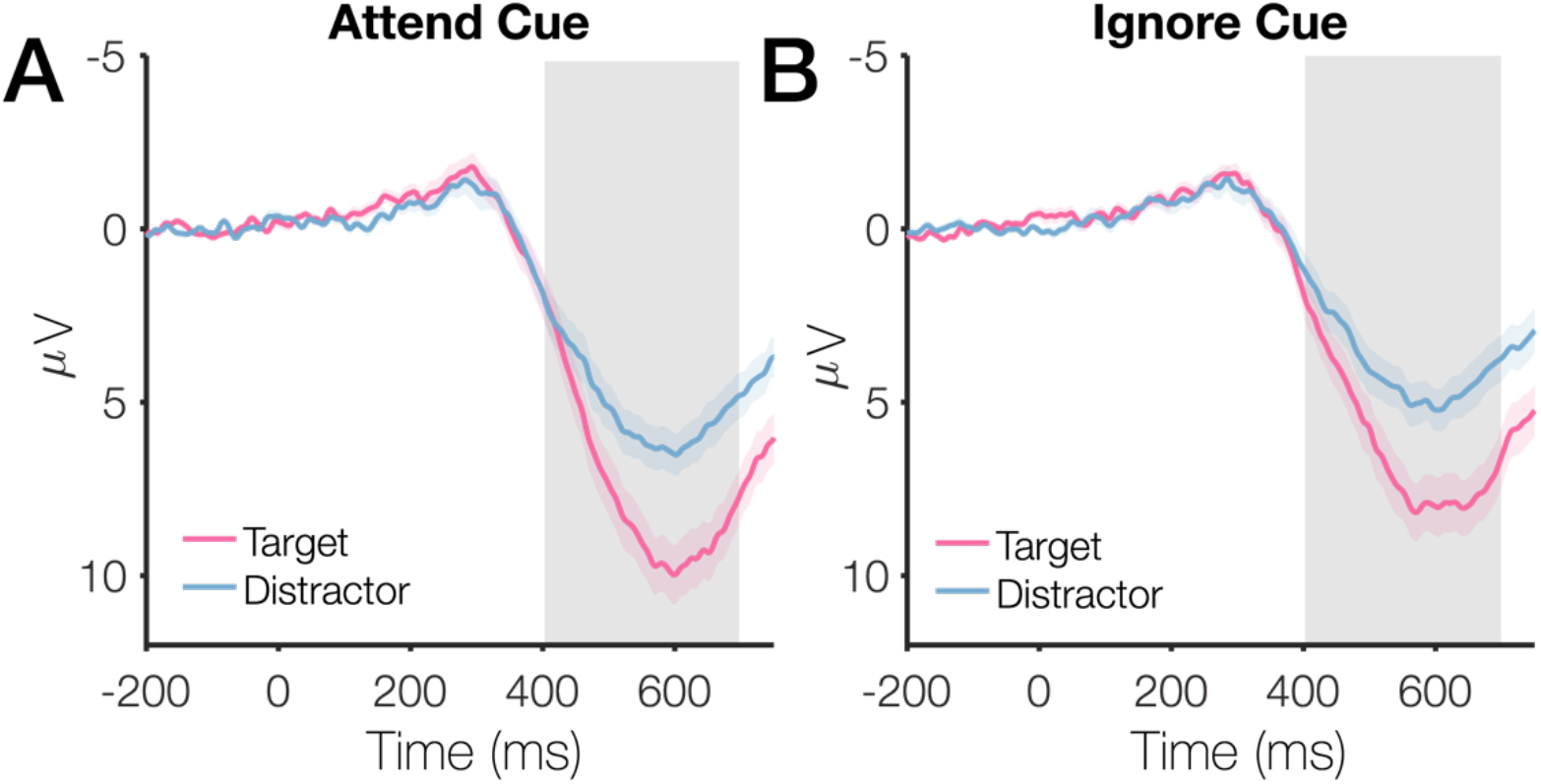
P3 amplitude at electrodes Pz and POz. (A) P3 amplitude in the attend cue condition, as a function of whether the event onset (iso-oriented lines) was a target that should be reported or a distractor that should be ignored. (B) P3 amplitude in the ignore cue condition. Shaded error bars represent standard error of the mean; the gray rectangle indicates the time period used for the statistical tests.

### Focused review of feature-based attention studies using SSVEPs

Given that our results are inconsistent with prior work, we conducted a focused review to try to pinpoint critical methodological differences that may have led to our failure to replicate a basic attention effect on SSVEP amplitude in this specific task. To do so, we first read review papers to identify an initial set of empirical studies employing a feature-based attention manipulation and SSVEPs (Andersen, Müller, et al., 2011; Norcia et al., 2015; Vialatte et al., 2010). From this initial set of papers, we used Google Scholar to check citations and citing papers for mention of the terms feature-based attention and SSVEPs. Our inclusion criteria included: (1) published journal article (2) healthy young adults (3) SSVEPs were measured from either an EEG or MEG signal and (4) a feature-based attention manipulation was included.

We defined “feature-based attention manipulation” as having the following characteristics: (1) Participants were cued to attend a feature(s) within a feature dimension (e.g., attend red, ignore blue) rather than across a feature dimension (e.g. attend contrast, ignore orientation), (2) The attended and ignored feature were both frequency-tagged in the same trials (rather than only 1 feature tagged per trial), (3) Each frequency was both “attended” and “ignored” on different trials, so that the amplitude of a given frequency could be examined as a function of attention, (4) The task could not be performed by adopting a strategy of splitting spatial attention to separate spatial locations.

After applying these screening criteria, some of the studies that we initially identified were excluded (brackets indicate exclusion reason(s)): Appelbaum & Norcia, 2009 [1,4]; Boylan et al., 2019 [3]; Bridwell & Srinivasan, 2012 [2,3]; Clementz et al., 2008 [3]; Garcia et al., 2013 [1,4]; Hasan et al., 2017 [2]; Itthipuripat et al., 2019 [1]; Talsma et al., 2006 [1,4]; Thigpen et al., 2019 [4]; Verghese et al., 2012 [1,4]).

We identified a total of 34 experiments from 28 unique papers (Appendices P-S) meeting our inclusion criteria. From these experiments, we quantified variables such as the number of subjects, number of trials, frequencies used, and the presence or absence of an attention effect in the expected direction (attended > ignored). If more than one group of participants was used (e.g., an older adults group) then we included the study but only quantified results for the healthy young adult group (Quigley et al., 2010; Quigley & Müller, 2014).

### Task used in each study

The tasks used in these studies fell broadly into one of 4 categories: (1) a competing gratings task, (2) a whole-field flicker task, (3) a hemifield flicker task and (4) a central task with peripheral flicker.

In the competing gratings task (Appendix P), participants viewed a stream of centrally-presented, superimposed gratings (e.g., a red horizontal grating and a green vertical grating). Because colored, oriented gratings were typically used, participants could thus generally choose to attend based on either one or both features (color and/or orientation). Each grating flickered at its own frequency (e.g. green grating shown at 7.41 Hz, red grating shown at 8.33 Hz, as in Chen et al., 2003). Because the gratings were superimposed, on any given frame only one of the two gratings was shown. On frames where both gratings should be presented according to their flicker frequencies, a hybrid “plaid” stimulus was shown. Studies using a competing gratings task include: (Allison et al., 2008; Chen et al., 2003; Keitel & Müller, 2016; J. Wang et al., 2007).

In the whole-field flicker task (Appendix Q), participants viewed a spatially global stimulus comprised of small, intermingled dots or lines. Typically, half of the dots or lines were presented in one feature (e.g., red) and the other half of the lines were presented in another (e.g., blue); each set of dots flickered at a unique frequency. Although the most common attended feature was color, some task variants included (1) attending high or low contrast stimuli (2) attending a particular orientation or (3) attending a particular conjunction of color and orientation. The whole-field flicker task was the most common task variant, and it is also most similar to the task performed here. Studies using a whole-field flicker task include: (Andersen et al., 2008, 2009, 2012, 2015; Andersen & Müller, 2010; Forschack et al., 2017; Martinovic et al., 2018; Martinovic & Andersen, 2018; Müller et al., 2006; Quigley et al., 2010; Quigley & Müller, 2014; Steinhauser & Andersen, 2019; D. Zhang et al., 2010)

In the hemifield flicker task, participants viewed a stimulus within each hemifield, and each stimulus was comprised of small, intermingled dots or lines. Often, these studies included both a feature-based attention manipulation and a spatial attention manipulation (e.g., attend red on the left-hand side). However, the feature-based attention task could not be achieved with spatial attention alone, as participants needed to attend to a particular color and ignore a distractor color within the attended hemifield. In addition, the unattended hemifield could often be used to track the spatially global spread of feature-based attention. Studies using a hemi-field flicker task include: (Adamian et al., 2019; Andersen et al., 2013; Andersen, Fuchs, et al., 2011; Müller et al., 2018; Störmer & Alvarez, 2014)

Finally, in the central task with peripheral flicker, participants performed a task near fixation (e.g., visual search), and feature-based attention was measured indirectly via a peripheral flickering stimulus (i.e., this design took advantage of the spatially global spread of feature-based attention, Sàenz et al., 2002, 2003; White & Carrasco, 2011). Studies using a central task with peripheral flicker include: (Chu & D’Zmura, 2019; Jiang et al., 2017; Painter et al., 2014, 2015).

### Sample Size, Trial Counts, and Stimulus Duration

First, we examined whether insufficient power could have led to our failure to detect an attention effect. Both sample size and the number of trials per condition are critical for determining power (Baker et al., 2019; Boudewyn et al., 2018; Button et al., 2013; Button & Munafò, 2017; Clayson et al., 2019; Thigpen et al., 2017). The number of studies employing each task variant is plotted in Figure 6A, the number of subjects per experiment is plotted in Figure 6B, the number of trials per experiment is plotted in Figure 6C, and stimulus duration is plotted in Figure 6D. Bars are color coded according to whether each experiment overall found an expected attention effect (attended > ignored), a reverse attention effect (ignored > attended), mixed results across conditions within the experiment, or ambiguous results (3 studies: Pei et al., 2002: reported statistics for the harmonics but not the fundamental frequency; Allison et al., 2008: missing formal statistical tests; Martinovic et al., 2018: statistical tests measured if attention effect differed between conditions, but did not formally test that the attention effect was overall significant). Our study had N = 23, 1,440 total trials per subject (720 trials per cue condition), and a stimulus duration of 2 seconds. In comparison to the literature, these factors are unlikely to explain our failure to find an effect. On average, prior studies had a median sample size of N = 15 (SD = 4.1, min = 9, max = 23) and a median trial count of 440 (SD = 314.2, min = 8, max =1600) and a median stimulus duration of 4.1 sec (SD = 37.35, min = 1 sec, max = 120 sec). Likewise, our trial counts were reasonable when we restricted our analysis to trials without targets or distractors (360 trials total; 180 trials per condition) relative to this estimated value for the literature (median = 96 trials per condition, SD = 95.6, min = 4, max = 400). Upon initially plotting the data, we noticed that studies using the competing gratings task produced inconsistent results, with as many experiments showing a mixture of effects across conditions, overall reversed effects (ignored > attended), and null effects (2 studies in each category). We think this inconsistency is most likely due to the below-average trial counts for these studies (median of only ~8 trials) (Boudewyn et al., 2018).

**Figure 6.**
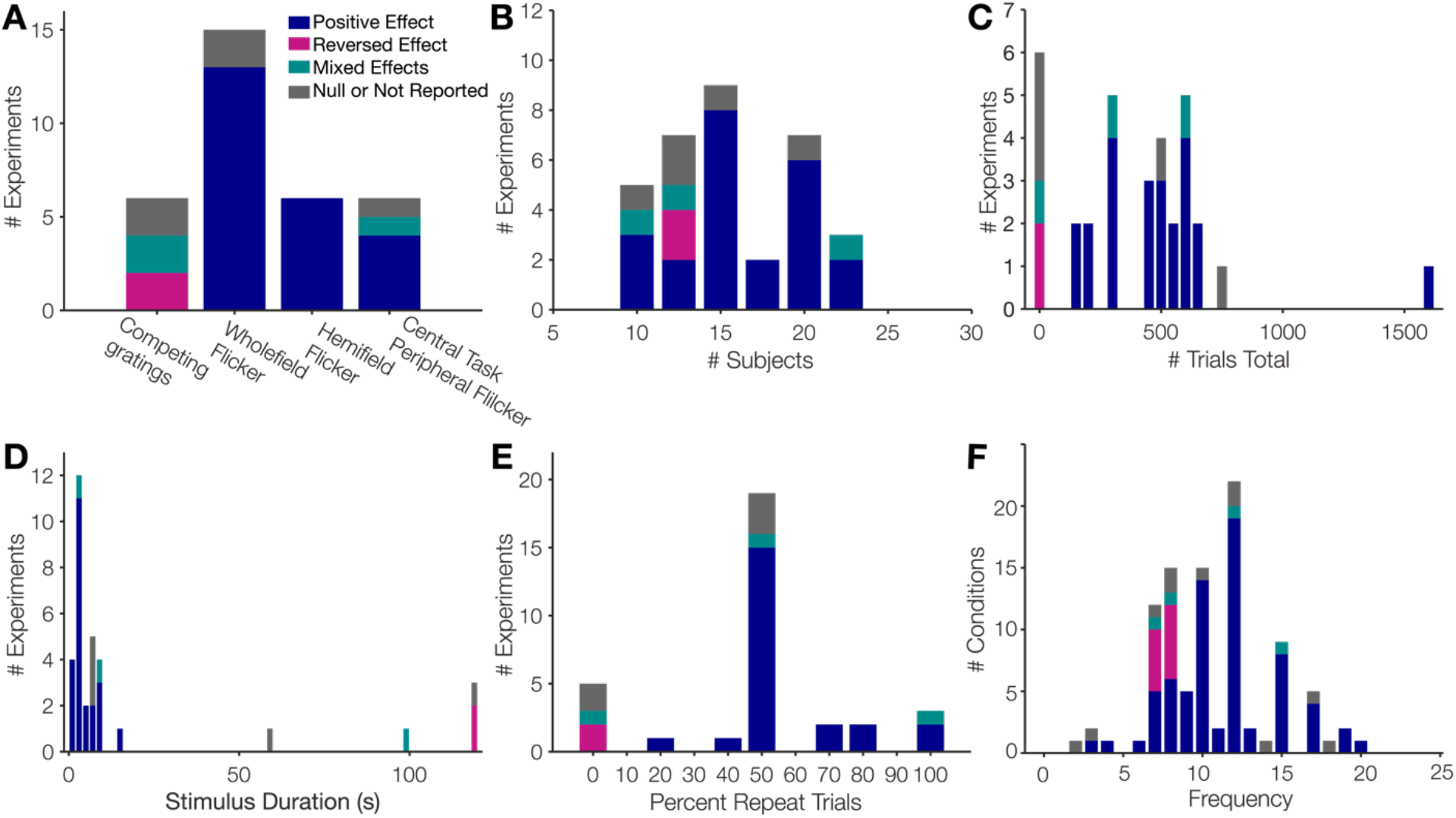
Study characteristics from the literature review of feature-based attention and SSVEPs. (a) Number of experiments in each of the 4 task variants. Different colors in the stacked bar graphs indicate whether the experiment found an expected attention effect (attended > ignored), reverse effect (ignored > attended), mixed effects across conditions, or a null / ambiguously reported effect. (b) Number of participants. (c) Number of trials in the experiment (before artifact rejection or excluding trials with targets). (d) Stimulus duration. (e) Percentage of trials, on average, where the attended feature was repeated on the next trial. (f) Stimulation frequencies.

### Percentage of trials where an attended feature was repeated

Next, we examined the percentage of trials where the attended feature was repeated (e.g., if the attended color was red on trial *n*, what was the chance that red would also be attended on trial *n*+1?). The priming-based account of feature-based attention posits that participants cannot use trial-by-trial cues to enhance a particular feature, but rather, feature-based enhancement happens automatically when a particular feature is repeated (Theeuwes, 2013). Thus, if there is a substantial proportion of trials where the repeated color was attended (e.g. with 2 possible colors, both the attended and ignored color will be repeated on 50% of trials), then the observed attentional enhancement effects might be driven primarily by incidental repetitions of attended features. In our study, we used 5 different colors to reduce the potential effect of inter-trial priming on the observed SSVEP attention effects (20% repeats of the attended color, 4% repeats of the attended color and the ignored color). We quantified the approximate percentage of trials on which an attended feature on one trial is repeated on the next trial (within a given block of trials). In some studies, participants were cued to attend more than one feature on a given trial, or they sometimes attended to a conjunction of features. In these cases, we calculated the expected number of repeats for either of the 2 attended features based on the total number of conditions (Andersen et al., 2008, 2013, 2015; Martinovic et al., 2018).

Figure 6E shows a histogram of the percentage of trials that repeated an attended feature. In two studies, the to-be-attended color was held constant on each block (100% repeats (Jiang et al., 2017; Painter et al., 2014). In the majority of the remaining studies, only two unique features were used so the percentage of trials where the attended feature was repeated was on average quite high (overall median = 50%, SD = 27.4%, min = 0%, max = 100%). Finally, in three studies, the attention conditions were perfectly alternated (0% repeat trials; Allison et al., 2008; Chen et al., 2003; J. Wang et al., 2007). Consistent with a priming account, 81% of the studies with a high percentage of repeats showed a consistent positive attention effect, whereas none of the studies with 0% repeat trials showed a consistent positive attention effect. However, we think the inconsistent effects in the studies with 0% repeats might be equally attributed to their low trial counts (median = 8 trials per condition; Boudewyn et al., 2018). Only one study had a similar proportion of repeats as the present study (Störmer & Alvarez, 2014). Störmer and Alvarez found a significant attention effect while using 5 unique colors (intermixed randomly from trial to trial). The findings by Störmer and Alvarez provide evidence against the feature-based priming account, and suggest the task factor “number of colors” cannot definitively explain our inability to observe an attention effect. However, given the lack of extant work using unpredictable color cues, we think future, systematic work is needed to determine the degree to which inter-trial priming effects may modulate the size and reliability of feature-based attention effects.

### SSVEP Frequencies

We examined frequencies that have been most commonly used in the literature. In our study, we chose relatively high frequencies (24 and 30 Hz) in order to have increased temporal resolution for detecting potential time-course effects. In addition, some have argued that using higher frequencies as advantages for driving a more localized portion of visual cortex, as opposed to broadly driving visual, parietal and frontal cortex when using lower frequencies in the theta/alpha bands (i.e., ~7-12Hz; Ding et al., 2006; Srinivasan et al., 2006). For the purposes of temporal resolution, earlier work examining the time course of *spatial* attention with SSVEPs used the frequencies 20 and 28 Hz, (Müller et al., 1998). However, upon reviewing the feature-based attention literature, we found that our chosen frequencies were outside the range that has previously been used with a feature-based attention task (Figure 6F; median = 10 Hz, SD = 3.6 Hz, min = 2.4 Hz, max = 19.75 Hz). Thus, it is possible that feature-based attention, unlike spatial attention, cannot be easily tracked with flicker frequencies above ~20 Hz. Although some studies have argued that feature-based attention effects do not qualitatively appear to vary with frequency (Martinovic et al., 2018; Steinhauser & Andersen, 2019), the vast majority of reviewed studies only reported statistical significance of an overall attention effect collapsed across frequencies. Only a handful studies have reported statistical significance of individual frequencies (e.g., Chu & D’Zmura, 2019; Painter et al., 2014; Quigley & Müller, 2014; Steinhauser & Andersen, 2019). As such, future work is needed to systematically investigate the effect of frequency choice on feature-based attention effects, particularly for frequencies >20 Hz.

### Task Type and Task Difficulty

Finally, we examined whether the type of task and task difficulty may have influenced our ability to detect an attention effect. In particular, the specific targets that we used may differ slightly from prior work. In our experiment, participants detected a brief period (333 ms) of an on average ~75% coherent orientation (the coherent line orientation was a random, unpredictable direction, from 1-180 degrees). In this task, participants performed well above chance, but the task was still fairly challenging overall (d’ = 1.25). This raises the possibility that, compared to prior SSVEP studies, subjects were giving up on some percentage of the trials and that this contributed to the lack of attention effects.

For the reviewed papers in which participants detected a target within the flickering stimulus (“whole-field flicker task” and “hemifield flicker task”), we compiled information about participants’ accuracy, the duration of the target, the type of target, and the percentage of dots/lines that comprised the target (Table S5). We found that our particular task (detect a coherent orientation in the cued color) was slightly different from the other tasks that have been used. Two other prior studies did not use a behavioral task at all: participants were simply instructed to monitor a particular feature without making any overt response (Pei et al., 2002; D. Zhang et al., 2010). In three papers, participants attended to a brief (200 ms) luminance decrement in 20% of the attended dots (Adamian et al., 2019; Andersen et al., 2009, 2013). In the remaining papers, participants monitored for a brief coherent motion event (230 ms – 500 ms) in 50-85% of the attended dots/lines (Andersen et al., 2008, 2012; Andersen & Müller, 2010; Forschack et al., 2017; Martinovic et al., 2018; Martinovic & Andersen, 2018; Müller et al., 2006, 2018; Quigley et al., 2010; Quigley & Müller, 2014; Steinhauser & Andersen, 2019; Störmer & Alvarez, 2014).

Although the particulars of the luminance and motion tasks subtly differ from our orientation task, it is not clear why SSVEPs would track attention when the target is a coherent luminance value or motion direction, but not when the target is a coherent orientation. For example, just like in the coherent motion direction tasks used by others, the angle of the coherent orientation in our task was completely unpredictable. Thus, participants in our task and in other tasks could not form a template of an orientation or motion direction they should attend in advance and instead had to attend to an orthogonal feature dimension such as color. In addition, in both prior tasks and the current task there were an equal number of coherent events in the cued and uncued color. If participants failed to attend to the cued color and instead responded to any orientation event, their performance in the task would be at chance.

Behavioral performance in the reviewed studies ranged from d’ = 0.8 to d’ = 3.25 (Appendix T). In many studies, performance was quite high (d’ > 2 or accuracy > 90%) relative to performance in our study (Adamian et al., 2019; Andersen et al., 2008, 2009, 2012, 2013; Forschack et al., 2017; Müller et al., 2006; Quigley et al., 2010; Quigley & Müller, 2014; Steinhauser & Andersen, 2019). However, there were several studies where the authors found SSVEP attention effects despite overall lower behavioral performance values more comparable to our study (d’ between 1 and 1.5; Andersen et al., 2015; Martinovic et al., 2018; Martinovic & Andersen, 2018). Sometimes, a more difficult task may actually be associated with increased attention effects: Martinovic and Andersen (2018) observed attention effects that were stronger in the subset of conditions with lower behavioral performance (d’ = .8 – 1.5) compared to conditions that were easier (d’ > 2.25).

## Discussion

In this pre-registered study, we sought to test whether cuing participants to *ignore* a particular color modulates the time-course of feature-based attention as indexed by steady-state visually evoked potentials (SSVEPs). As a baseline point of comparison, we also included a condition in which participants were cued to *attend* a particular color. This “attend cue” condition was intended as a close replication of much prior work showing that SSVEP amplitudes are modulated by attention (greater amplitude for the attended feature; e.g., Andersen et al., 2008; Andersen & Müller, 2010; Müller et al., 2006). However, we failed to replicate this basic overall attention effect; we found no difference in SSVEP amplitude as a function of attention in either the attend cue or the ignore cue condition. Thus, because we found no overall SSVEP attention effect, we were unable to test our hypotheses about how this attention effect was modulated by being cued to attend versus cued to ignore. Despite the lack of an SSVEP attention effect, positive control analyses indicated that that participants did successfully select the cued target color (i.e., we observed a significantly larger P3 component for target events in the attended color than in the ignored color).

Given our failure to observe an effect of attention on SSVEP amplitude with our task procedures, we performed a focused review of the literature to quantify key methodological aspects of prior studies using SSVEPs to study feature-based attention. Based on this review, we concluded that sample size and trial counts likely did not explain our failure to find an effect; our sample size and trial counts were near the maximum values found in the surveyed literature. Likewise, the range of accuracy values found in the literature suggests that task difficulty does not explain our failure to find an attention effect. However, two key, intentional design differences may have hampered our ability to find an effect: (1) the number of colors in our stimulus set and (2) the frequencies used to generate the SSVEP.

The first key design difference in our study was the number colors in our stimulus set. We purposefully minimized the influence of inter-trial priming on our estimates of feature-based attention (Theeuwes, 2013) by using 5 unique colors and randomly choosing target and distractor colors on each trial. According to a priming account of feature-based attention, a relatively high proportion of trials where the attended color is repeated back-to-back could inflate or even entirely drive apparent feature-based attention effects. Using 5 colors somewhat protects against this possibility, because it ensures that the attended color is repeated on 20% of trials, and both the attended/ignored colors are repeated on only 4% of trials. In the literature, we found that most studies had back-to-back color repeats on at least 50% of trials. It is thus plausible that inter-trial priming could contribute to observed attention differences in these studies. Contrary to a priming account, however, one study found robust feature-based attention effects using a set of 5 unique colors (Störmer & Alvarez, 2014), suggesting that participants can use a cue to direct feature-based attention even when the proportion of repeated trials is relatively low. To date, however, no study has directly manipulated the proportion of repeated trials or the number of possible stimulus colors in an SSVEP study. Given emerging evidence that history-driven effects play an important role in shaping both spatial and feature-based attentional selection (Adam & Serences, 2020; Awh et al., 2012; Failing et al., 2019; Geng et al., 2019; Kadel et al., 2017; B. Wang & Theeuwes, 2018a, 2018b; B.-Y. Won & Geng, 2020), we think that future work is needed to directly investigate whether and to what degree SSVEP estimates of feature-based attention are modulated by inter-trial priming.

The second key design difference in our study was the chosen set of frequencies. To ensure adequate temporal resolution to characterize time-course effects, we chose to use slightly higher frequencies (24 and 30 Hz). We believed these values would be reasonable, because an initial study of the time-course of spatial attention used SSVEP frequencies in a similar range (20 and 28 Hz; Müller et al., 1998). In addition, frequencies in the beta band (~15-30 Hz) have commonly been used in other SSVEP studies of spatial attention (Garcia et al., 2013; Kashiwase et al., 2012; Müller, Picton, et al., 1998; Müller & Hillyard, 2000; Toffanin et al., 2009; D.-O. Won et al., 2016), and SSVEPs are overall robust using a wide array of frequencies (at least 1 to 50 Hz; Herrmann, 2001; Zhu et al., 2010). However, some spatial attention studies have found no attentional modulation of SSVEPs in the beta band (Antonov et al., 2020; Gulbinaite et al., 2019), or have found effects only for the second harmonic of beta band frequencies (Garcia et al., 2013; Kim et al., 2007; Vissers et al., 2017). Further, the SSVEP amplitude, estimated spatial extent of the SSVEP signal, and the size of spatial attention effects vary with frequency (Ding et al., 2006; Gulbinaite et al., 2019; Herrmann, 2001). Given differences in the cortical processing of locations and features (M. R. Cohen & Maunsell, 2011; Haxby et al., 1994; Kastner & Ungerleider, 2000; Mishkin & Ungerleider, 1982; Owen et al., 1996), and differences in SSVEP spatial extent and strength with frequency (Ding et al., 2006; Gulbinaite et al., 2019; Lithari et al., 2016), it is plausible that feature-based attention can only be tracked with a limited range of frequencies (e.g., frequencies near the alpha band). Future work will be needed to systematically investigate the effect of SSVEP frequency on feature-based attention.

It is perhaps puzzling that frequencies above 20 Hz have been commonly used in the spatial attention literature but have not been used in the feature-based attention literature. The truncation of the frequency distribution in the reviewed literature could be a piece of the “file drawer” in action. It is possible that other researchers likewise discovered that they were unable to track feature-based attention using certain frequencies, but that these null results were never published due to journals’ and authors’ biases toward publishing positive results (Cooper et al., 1997; Dickersin, 1990; Dickersin et al., 1992; Dwan et al., 2008; Ferguson & Heene, 2012; Franco et al., 2014; Rosenthal, 1979) and biases against publishing negative results (i.e., “censoring of null results”, Guan & Vandekerckhove, 2016; Sterling, 1959; Sterling et al., 1995). Thus, our results highlight the practical and theoretical importance of regularly publishing null results. On the practical side, if prior null results had been published, we may have better known which frequencies to use or avoid, and we would have been able to test our key hypotheses. On the theoretical side, our results highlight how seemingly unimportant null results can have implications for theory when viewed in the context of the broader literature. For example, if certain frequencies track spatial but not feature-based attention, this may inform our understanding of the brain networks and cognitive processes differentially modulated by flicker frequency (Ding et al., 2006; Srinivasan et al., 2006).

In short, we found no evidence that SSVEPs track the deployment of feature-based attention with our procedures, and future methodological work is needed to determine constraints on generalizability of the SSVEP method for tracking feature-based attention. We performed a focused review of prior studies using SSVEPs to study feature-based attention, and from this review we identified two key factors (frequencies used; likelihood of inter-trial feature priming) that should be systematically investigated in future work.

## Contributions

K.A. & J.S. planned the study. K.A., L.C., and N.R. collected data and performed the literature review. K.A. performed analyses and drafted the manuscript. All authors revised the manuscript.

## Pre-registration information

We published a pre-registered research plan on the Open Science Framework prior to data collection (https://osf.io/kfg9h/).

## Data availability

Data and analysis code will be made available online on the Open Science Framework at https://osf.io/ew7dv/ upon acceptance for publication.

## Acknowledgements

We thank Matteo d’Amico for additional assistance with data collection and Kelvin Lam for help as lab manager. We thank Rosanne Rademaker and Angus Chapman for sharing Psychtoolbox code that was adapted for use here.

---

**Appendix A.**
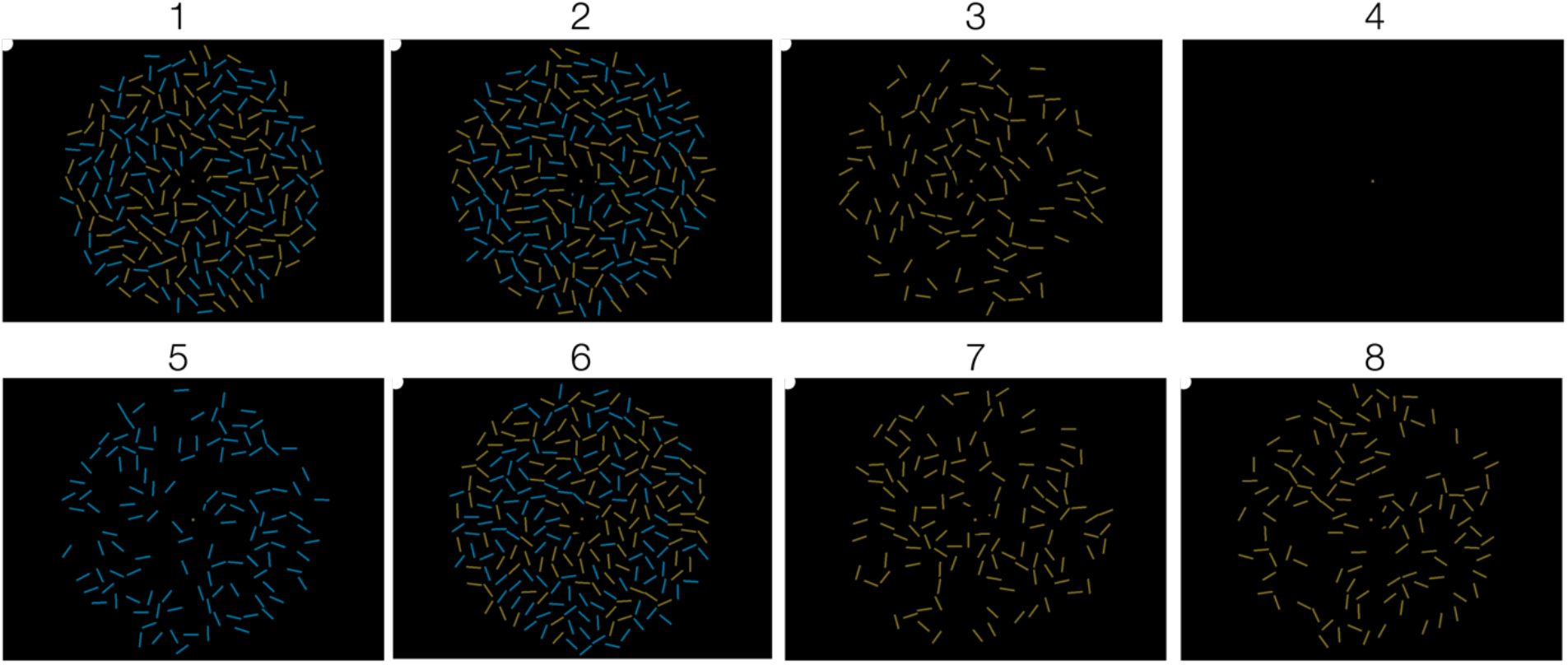
Example frames during the stimulus presentation. Eight example frames (1-8) from the stimulus presentation period illustrate how the flicker was achieved (refresh rate was 120 Hz, so each frame was ~8.33 ms). In this example, the attended color is yellow, and the attended frequency is 24 Hz (3 frames on, 2 frames off). Blue is the unattended color (30 Hz; 2 frames on, 2 frames off). The white dot in the upper left-hand corner was used to record the attended frequency flicker using a photodiode (this corner of the screen was covered with thick, opaque black electrical tape so that it was not visible to the participants.

**Appendix B.**
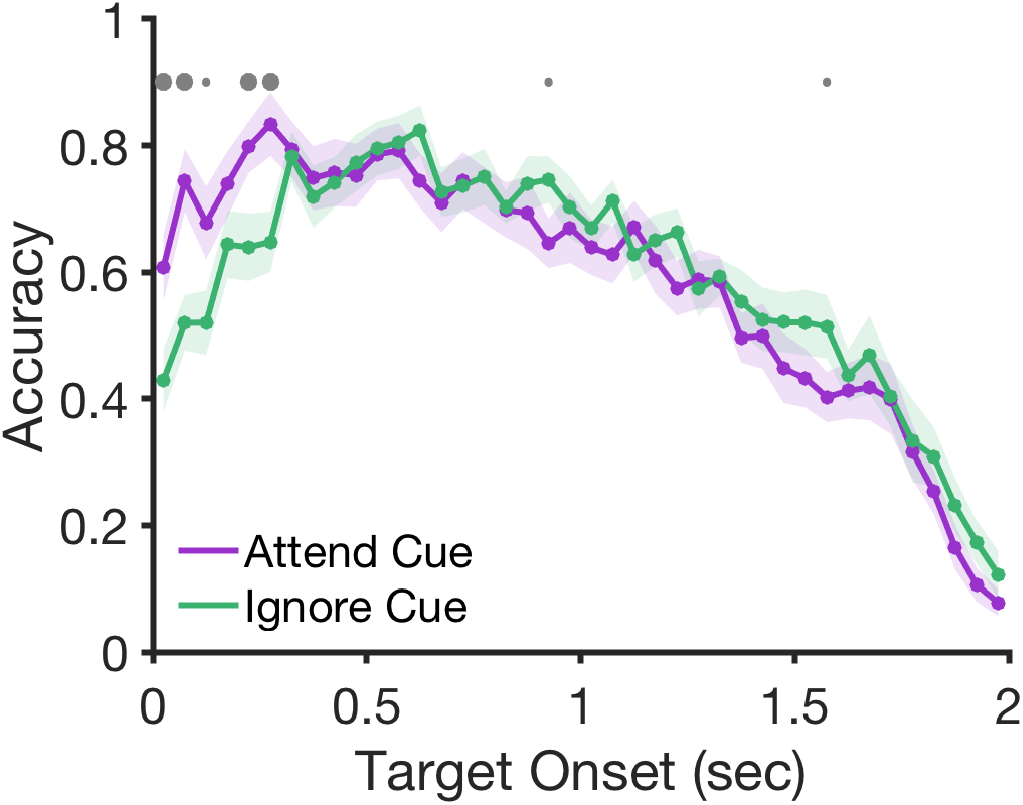
Accuracy for target-present trials as a function of the time between Cue Onset and the Target Onset. For short cue-target intervals (<= 275 ms), participants were more accurate for attend cues than ignore cues. This pattern suggests that participants were more quickly able to utilize the attend cue than the ignore cue. Shaded error bars indicate +/- 1 SEM. Small gray dots indicate p < .05 (uncorrected), large dots indicate p < .001 (uncorrected).

**Appendix C.**
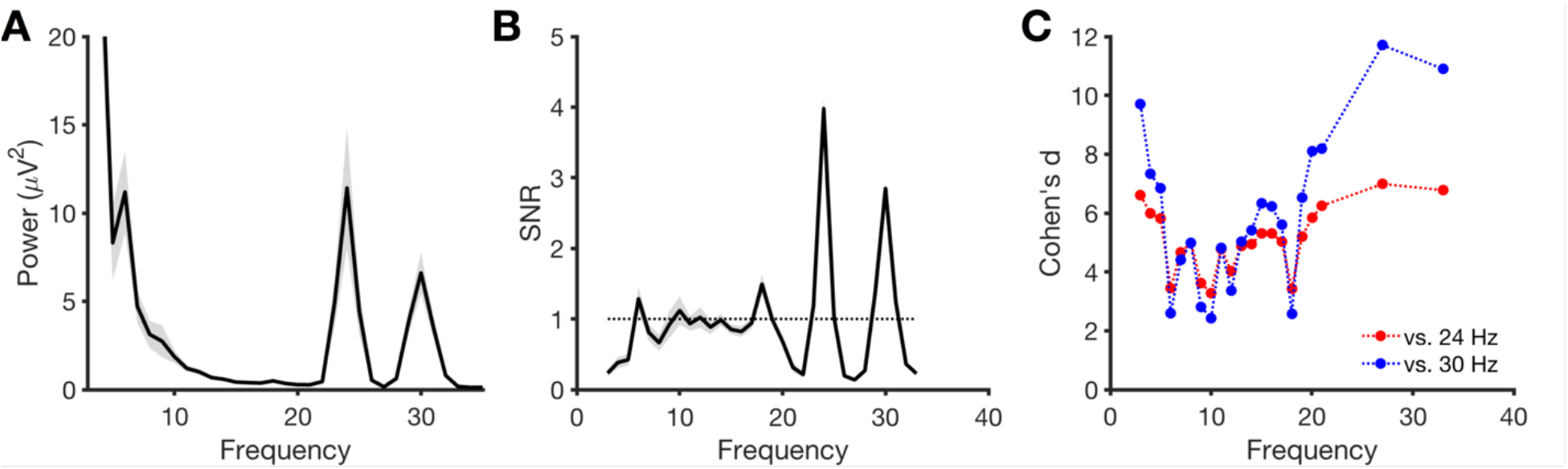
Power and SNR for each frequency. (A) Power for each frequency using the Gaussian wavelet filter analysis. (B) SNR for each frequency, calculated as the power at the frequency (e.g., 24 Hz) divided by the power at the average of the 2 neighboring 1-Hz frequencies on either side (e.g., average of 22, 23, 25, and 26 Hz). The theoretical chance level for SNR is 1 (dotted line), but because SNR is calculated with neighboring frequencies, frequencies that are adjacent to a significant “peak” may have values below 1. (C) Cohen’s d for the comparison between SNR at each of the two target SSVEP frequencies (24 Hz, 30 Hz) relative to other baselined frequencies (3-33 Hz excluding frequencies within +/- 2 Hz of the target SSVEP frequencies).

**Appendix D.**
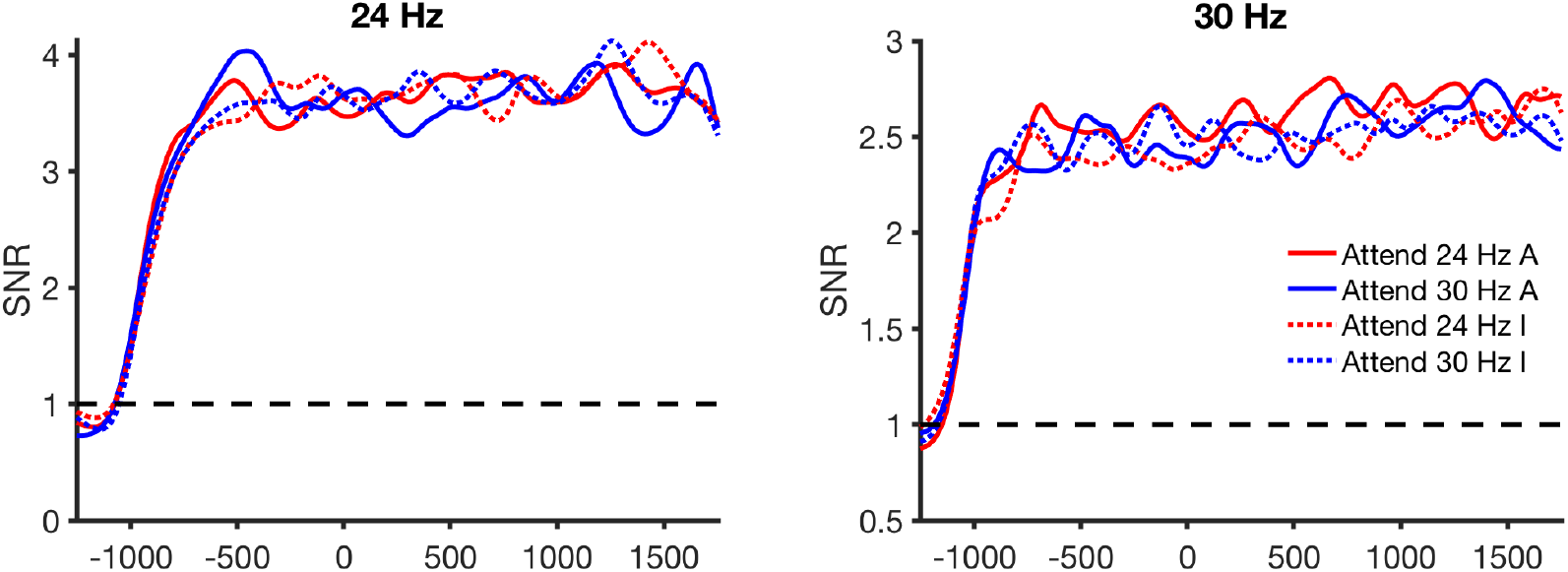
Time-course of SNR for each frequency. The stimulus began flickering at −1,333 ms, and the cue indicating which color to attend appeared at 0 ms. Red lines show when 24 Hz was the attended frequency; Blue lines show when 30 Hz was the attended frequency. Solid lines show data from the “attend cue” condition; Dotted lines show the “ignore cue” condition.

**Appendix E.**
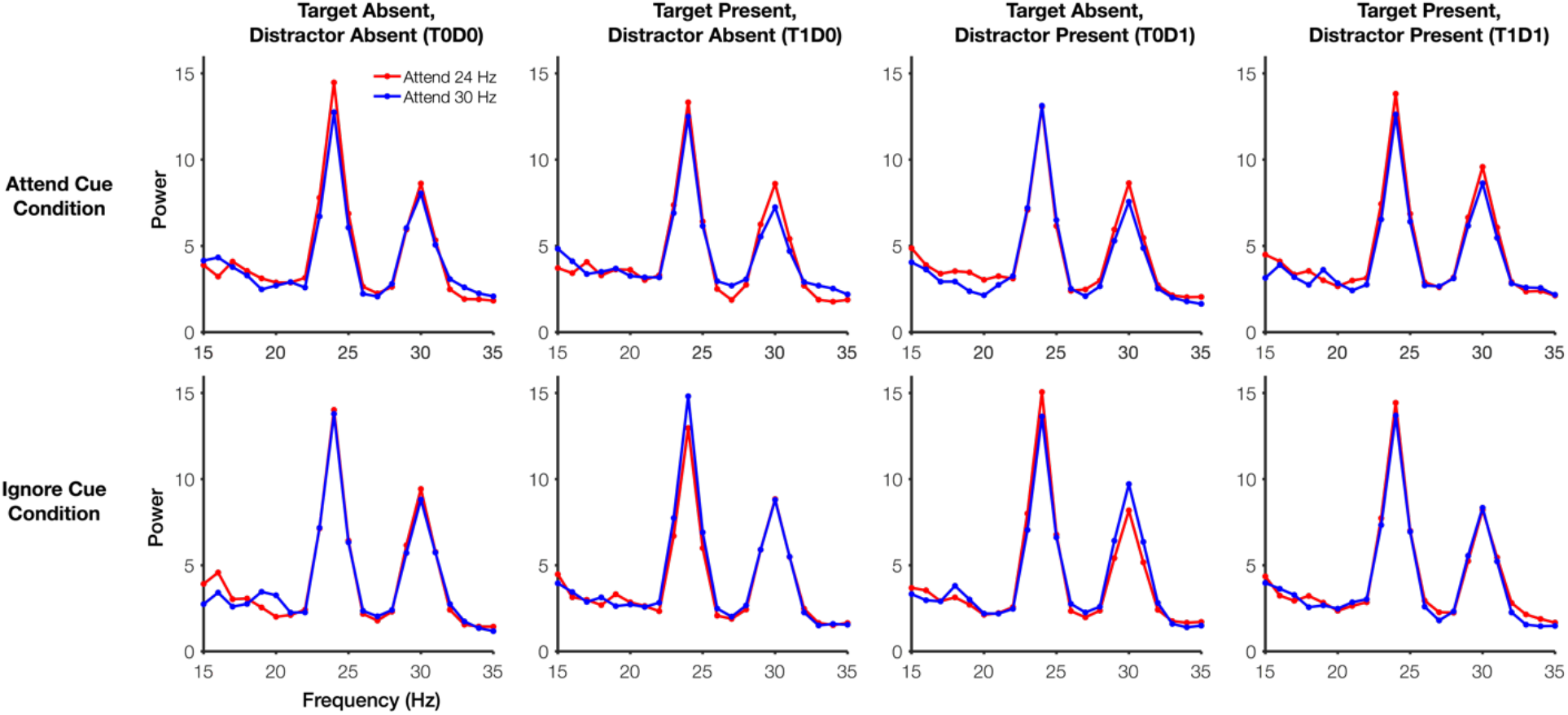
Frequency spectra separately for each target/distractor presence condition. Trials were counterbalanced to have a 50% chance of having a target event (T1) and to have 50% chance of including a distractor event (D1). Thus, 25% of trials had neither a target nor distractor (T0D0), 25% of trials had a target only (T1D0), 25% of trials had a distractor only (T0D1), and 25% of trials had both a target and a distractor (T1D1). Frequency spectra for each sub-condition are shown (Rows: Attend Cue or Ignore Cue, Columns: Each combination of target and distractor present/absent).

**Appendix F.**
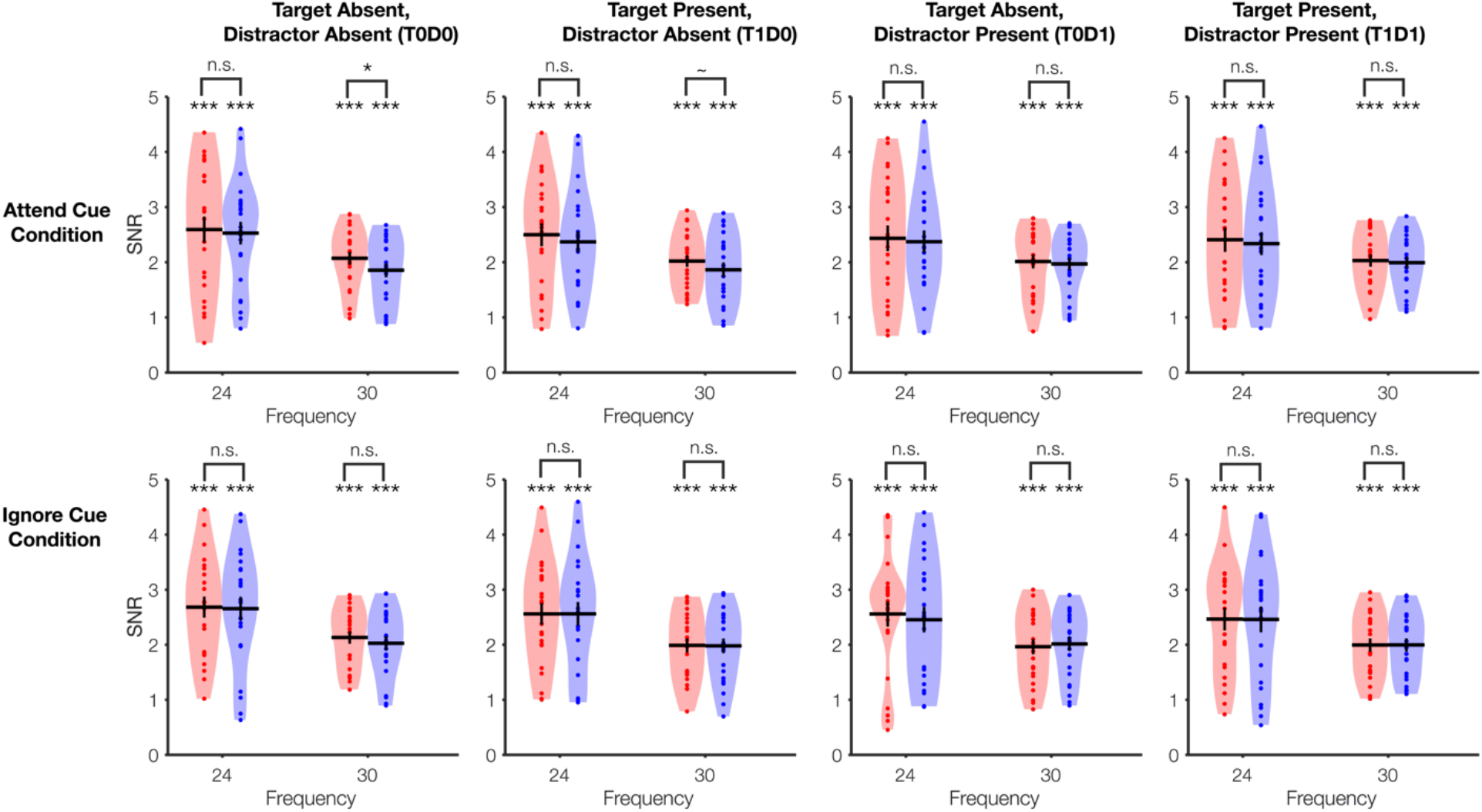
Signal to noise ratio (SNR) values separately for each target/distractor presence condition. Trials were counterbalanced have a 50% chance of having a target event (T1) and to have 50% chance of including a distractor event (D1). Thus, 25% of trials had neither a target nor distractor (T0D0), 25% of trials had a target only (T1D0), 25% of trials had a distractor only (T0D1), and 25% of trials had both a target and a distractor (T1D1). Frequency spectra for each sub-condition are shown (Rows: Attend Cue or Ignore Cue, Columns: Each combination of target and distractor present/absent). The bottom row of asterisks shows post-hoc, uncorrected significance for overall SSVEP signal compared to a null value of 1. The SSVEP signal was overall highly significant (***, p<.001). The top row of asterisks shows post-hoc, uncorrected significance for the comparison between the two adjacent bars (n.s. p > .10, ~ p <.10, * p < .05). Note, no conditions showed an attention effect (attended frequency > ignored frequency); the only significant, uncorrected post-hoc comparison was in the wrong direction (ignored > attended).

**Appendix G.**
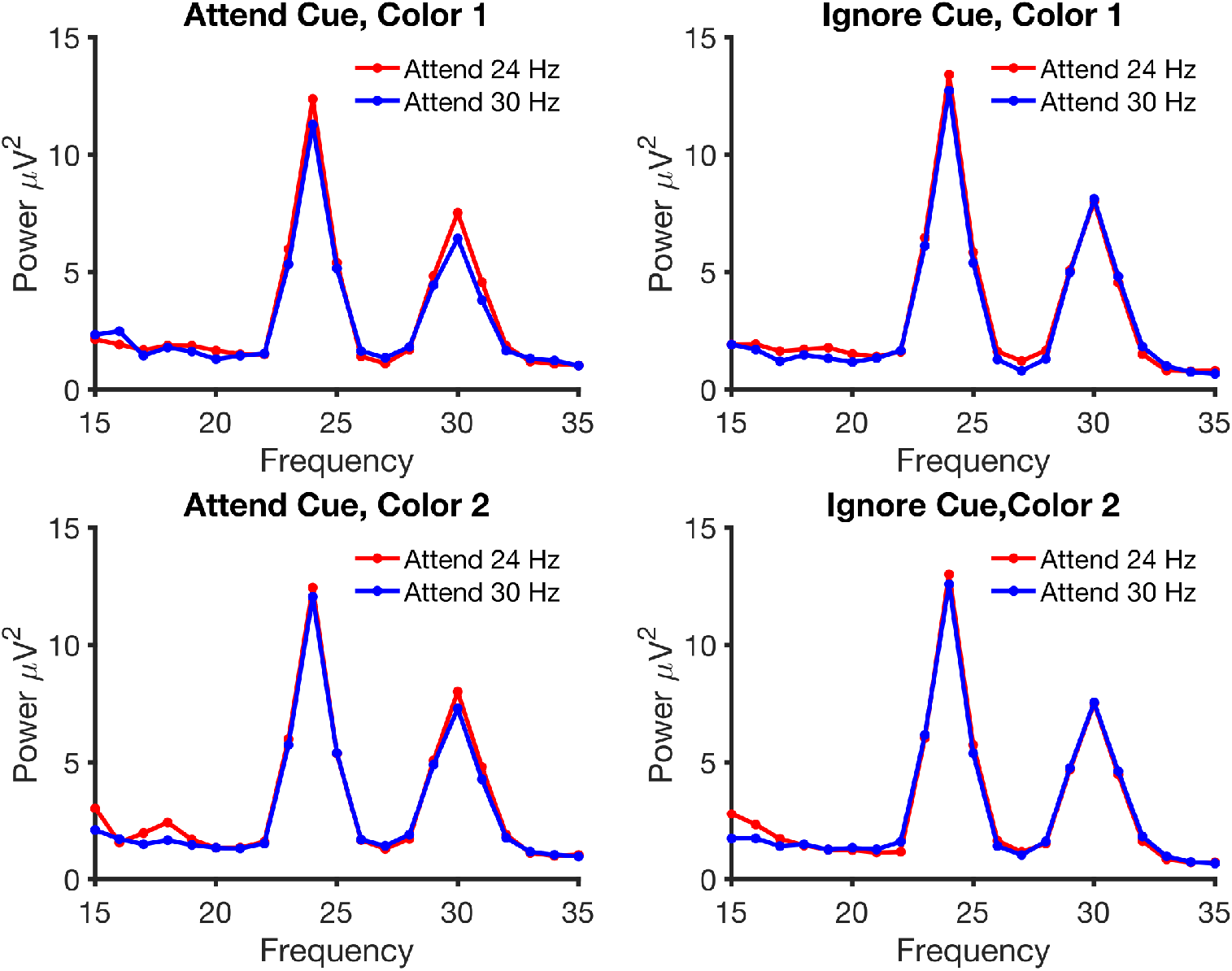
Power by frequency separately for each color distance condition. Target and distractor colors were randomly assigned on each trial from a pool of 5 possible colors. Thus, the target and distractor colors could be either 72 degrees (Color 1) or 144 degrees (Color 2) apart on a color wheel. We found no evidence of an attention effect in either color distance condition.

**Appendix H.**
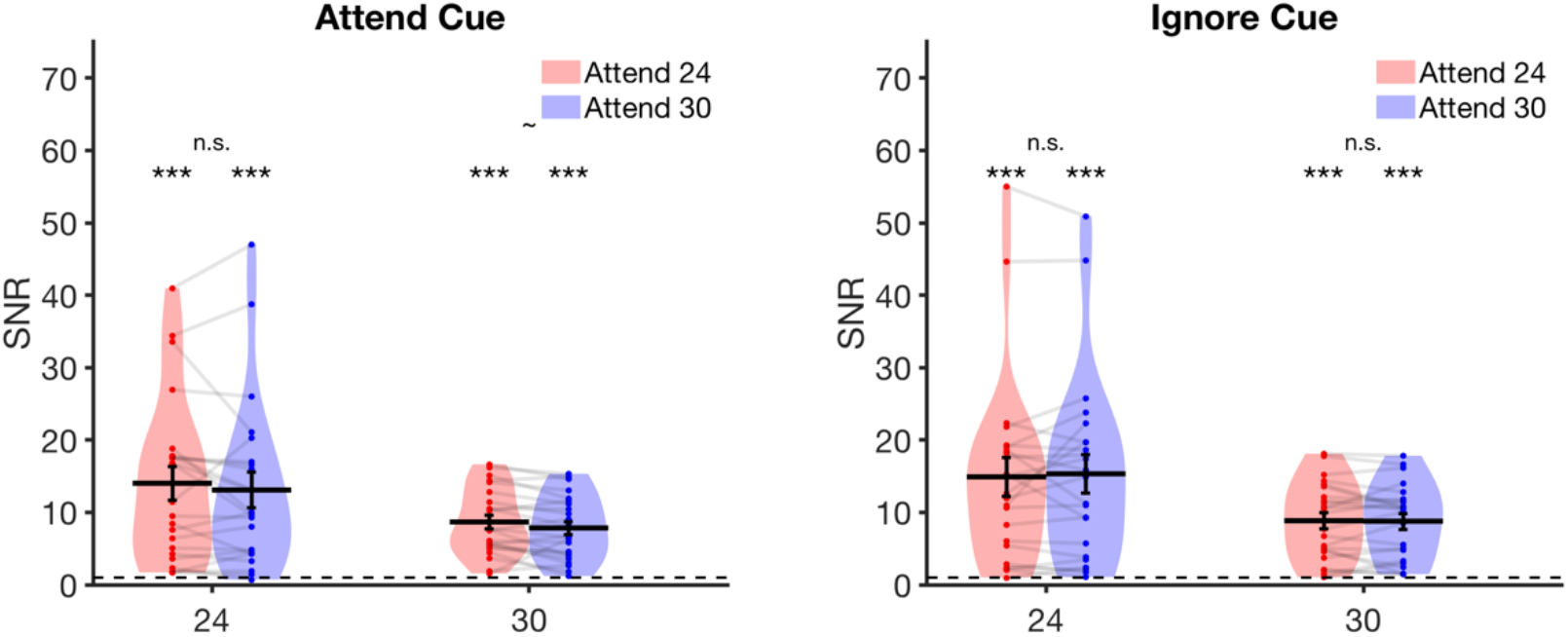
An additional analysis variant for the main SNR measure: skipping the first bin for computing SNR. Rather than using the pre-registered frequencies of +/- 1 and +/- 2 Hz for computing SNR, we instead skipped the first 1 Hz bin. Since +/-1 Hz had greater than baseline power, we may have attenuated our ability to observe SSVEP-related differences by including this bin in our SNR subtraction. For this analysis variant, we instead calculated SNR as the peak frequency minus the average of all frequencies +/- 2 and +/- 3 Hz from the peak (e.g., to compute SNR for 24 Hz, we subtracted the mean power at 21, 22, 26, and 27 Hz). Although overall SNR was much higher across all conditions using this metric, the pattern across experimental conditions was unchanged (i.e., we found no significant attention effects).

**Appendix I.**
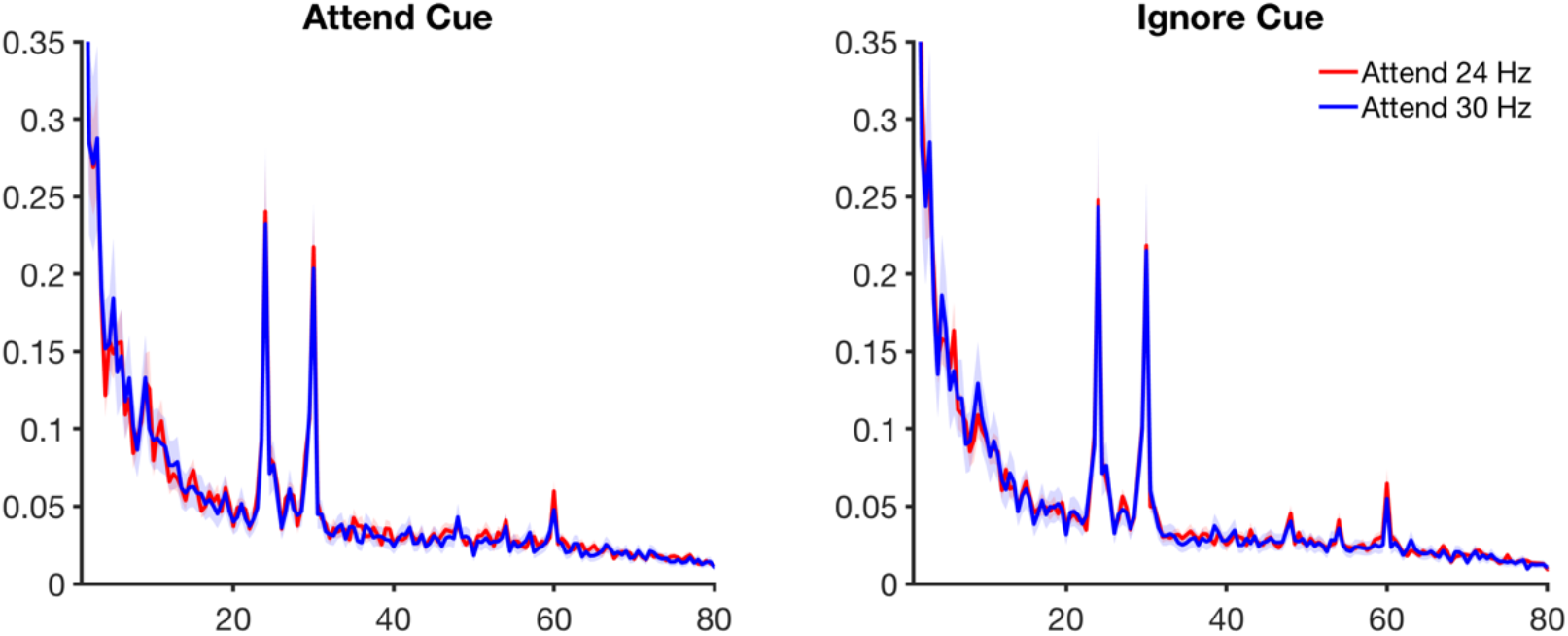
FFT analysis with a wider x-axis to show both the fundamental and second harmonic frequencies. (Left) FFT for the ‘attend cue’ condition. (Right) FFT for the ‘ignore cue’ condition. X-axis values are frequency (Hz); Y-axis values are amplitude (microvolts).

**Appendix J.**
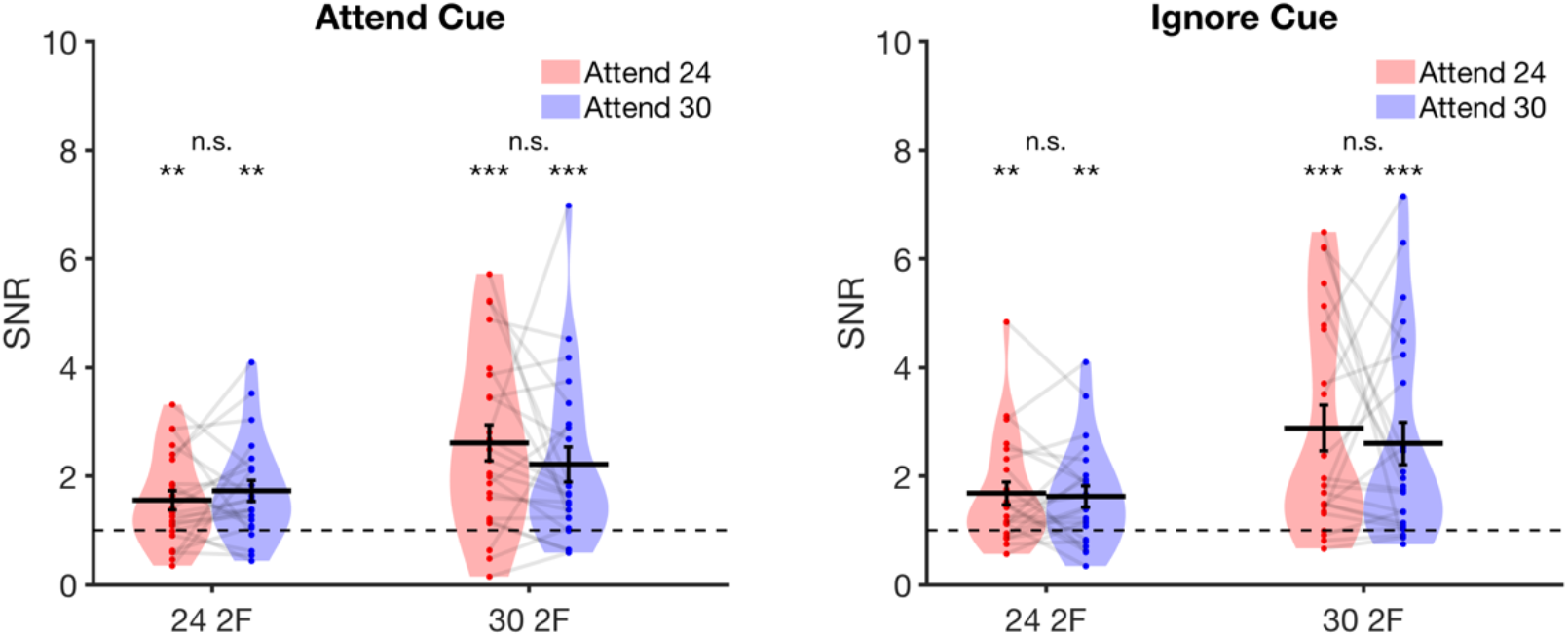
Violin plot of the second harmonic frequencies 48 Hz and 60 Hz from the FFT analysis. (Left) Violin plot of SNR for the second harmonic frequencies in the ‘attend cue’ condition; SNR for both harmonics was greater than 1, but there were no attention effects. (B) Violin plot of SNR for the second harmonic frequencies in the ‘ignore cue’ condition; SNR for both harmonics was greater than 1, but there were no attention effects.

**Appendix K.**
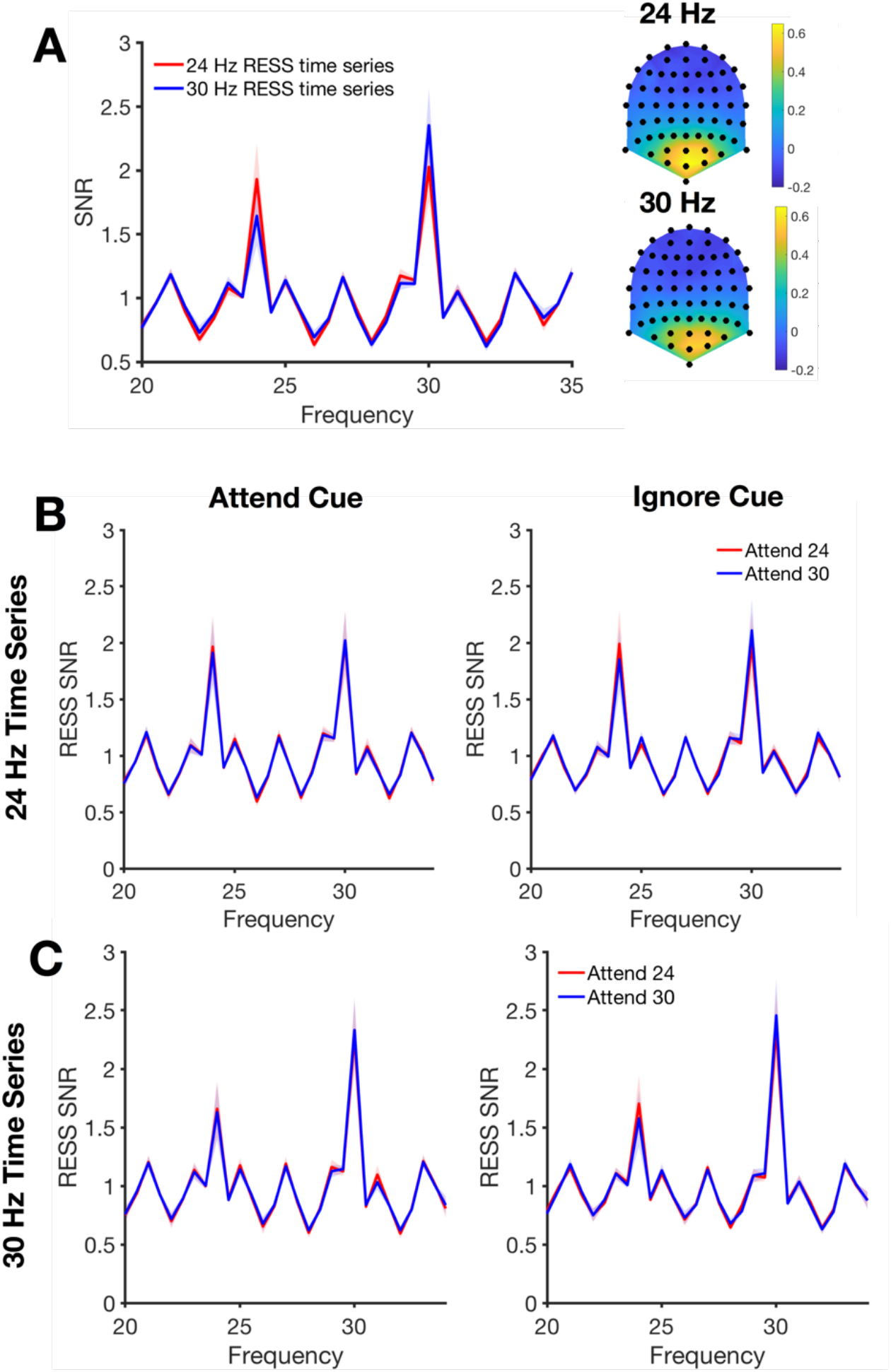
Rhythmic Entrainment Source Separation (RESS) analysis likewise shows null attention effects. Following code associated with [1], we performed rhythmic entrainment source separation (RESS) on our data to ensure that our a priori choice of electrodes did not impede our ability to find an attention effect. We decided to stick very closely to the default settings for RESS code developed by others in order to take some ‘researcher degrees of freedom’ out of the equation. We obtained a highly consistent pattern of results despite using a data-driven, single-trial approach that differs substantially from our pre-registered trial-averaged approach. We also note that the SNR values from the RESS approach are lower than the trial-averaged FFT we present in the main analysis, but that RESS does still provide an SNR advantage when compared to a single-trial FFT approach, as in [1]. We first calculated the spatial filters using data from all trials and the full trial length (−1000 ms to 2000 ms). We then applied the spatial filters to calculate SNR for each condition of interest (e.g., “Attend 24 Hz, Attend Cue Condition”, 24 Hz RESS time series; 500 ms to 2000 ms). For the analysis, we used a frequency resolution of 0.5 Hz, a full-width half maximum (FWHM) of 0.5 Hz for the center frequency, a FWHM of 1 Hz for the neighboring baseline frequencies +/- 2 Hz from the peak frequency. SNR was calculated as the ratio between each frequency of interest and the frequencies +/- 2 Hz away. (A) Normalized SNR by frequency and topography of the RESS time series optimized for 24 Hz (red) and 30 Hz (blue), collapsed across all conditions. (B) SSVEP response (computed as normalized SNR) for the 24 Hz-optimized RESS time series in the attend cue condition and ignore cue condition. (C) SSVEP response (computed as normalized SNR) for the 30 Hz-optimized RESS time series in the attend cue condition and ignore cue condition. We again found no significant effects of attention for either SSVEP frequency.’

**Appendix L.**
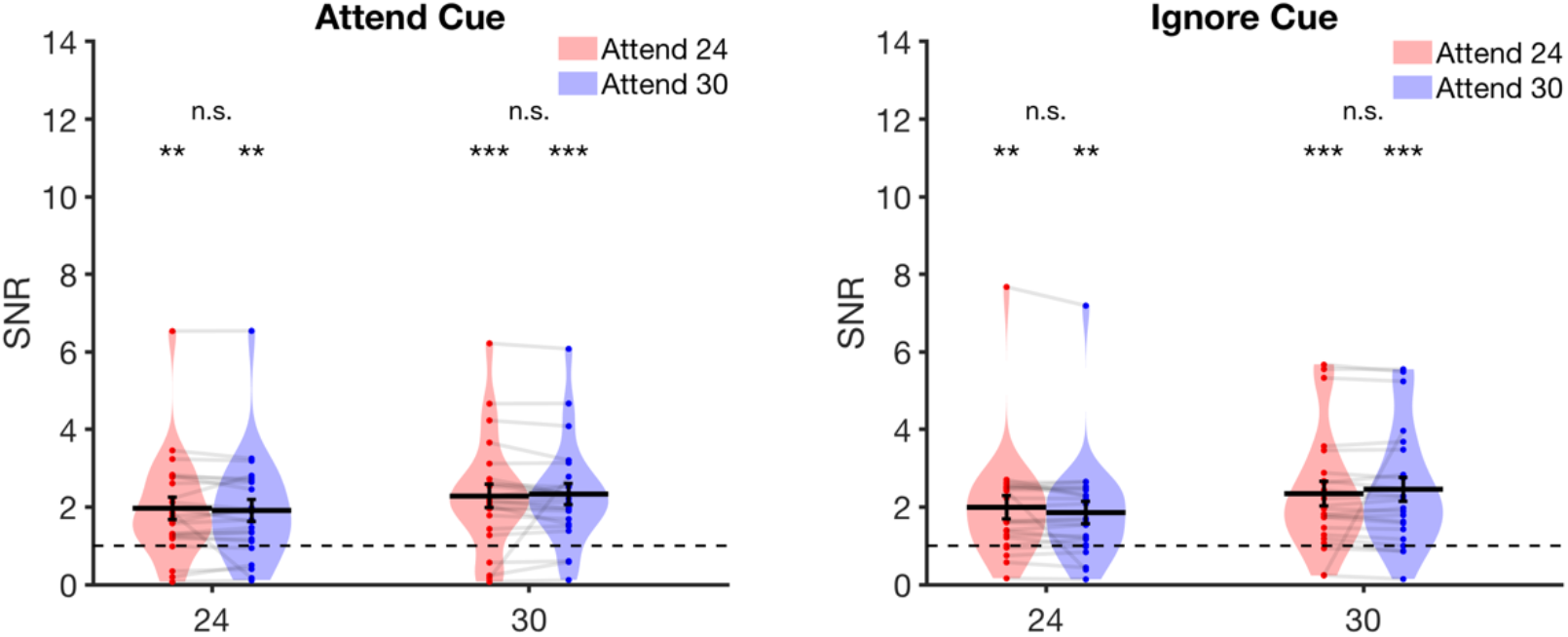
Violin plots of values obtained from the Rhythmic Entrainment Source Separation (RESS) analysis. We found no effect of attention on RESS values in either the Attend Cue condition (left panel) or the Ignore Cue condition (right panel).

**Appendix M.**
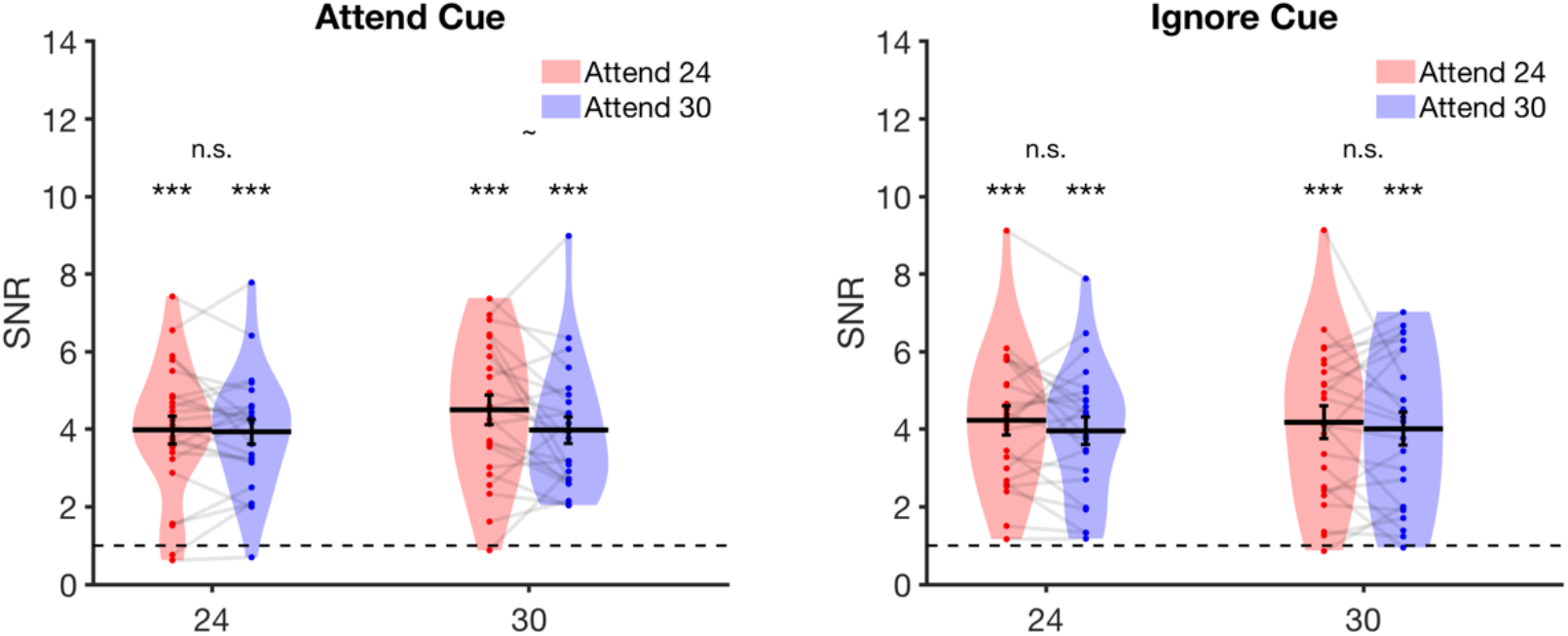
Violin plots of SNR values for each frequency, calculated from an FFT analysis on accurate trials only. Performing an FFT analysis on accurate trials only likewise yields null attention effects both in the attend cue condition (left panel) and the ignore cue condition (right panel).

**Appendix N.**
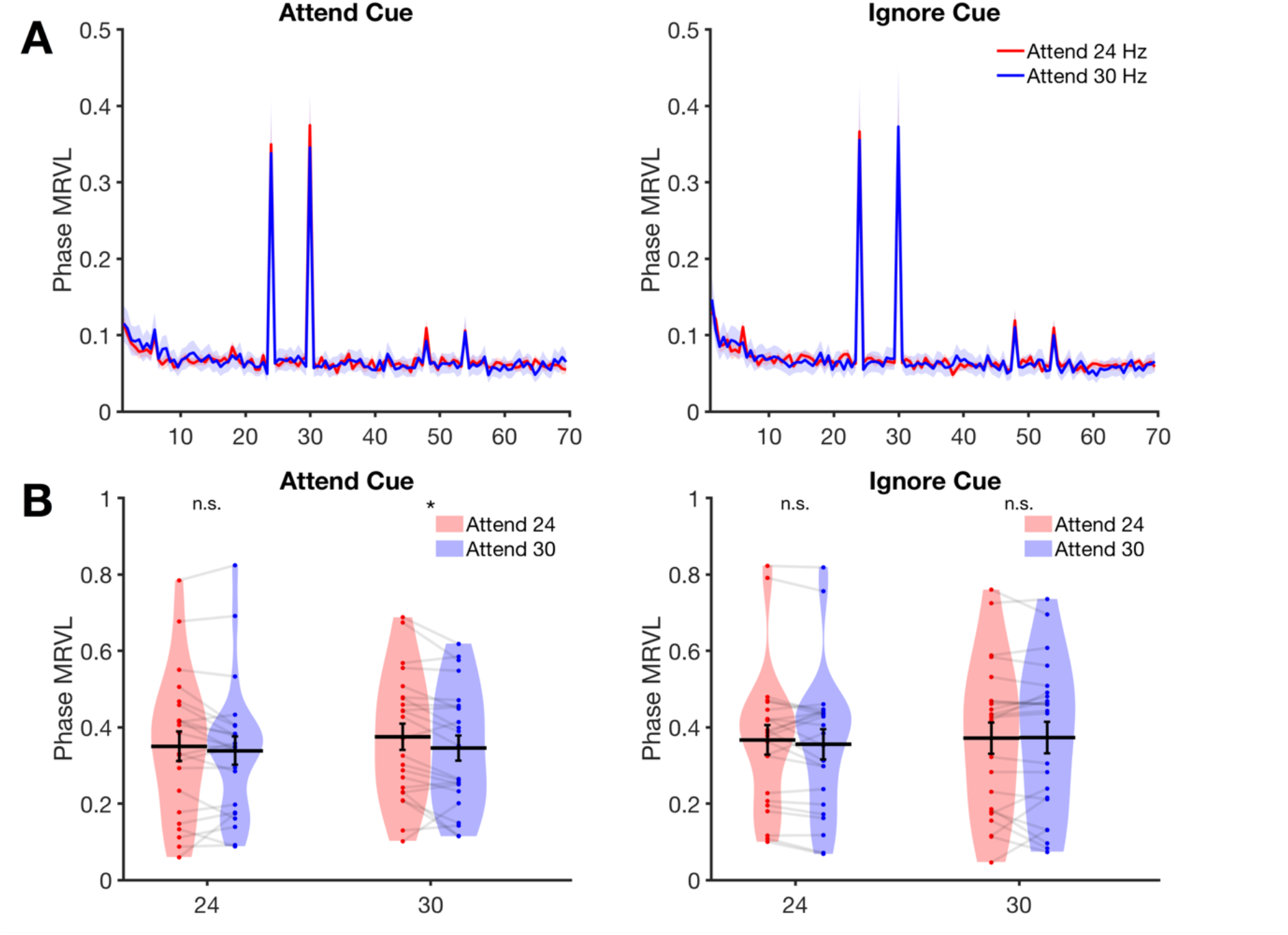
Results of the phase-locking index (PLI) analysis. We performed an FFT on single trials rather than on condition-averaged waveforms (time window: 333 ms – 2000 ms), and we extracted single-trial phase values (‘*angle.m*’). We calculated a phase-locking index by computing mean-resultant vector length on histograms of single-trial phase values (separate histograms for each condition, electrode, and frequency). Mean-resultant vector length ranges from 0 (fully random values) to 1 (perfectly identical values), for reference, see: Zar (2010). **(A)** Phase locking index (PLI) as indexed by mean-resultant vector length, averaged across electrodes O1, O2, and Oz. Replicating prior work, we found robust PLI values at the two SSVEP frequencies (24 and 30 Hz). **(B)** However, we found no evidence that PLI values were modulated by attention in the expected direction.

**Appendix O.**
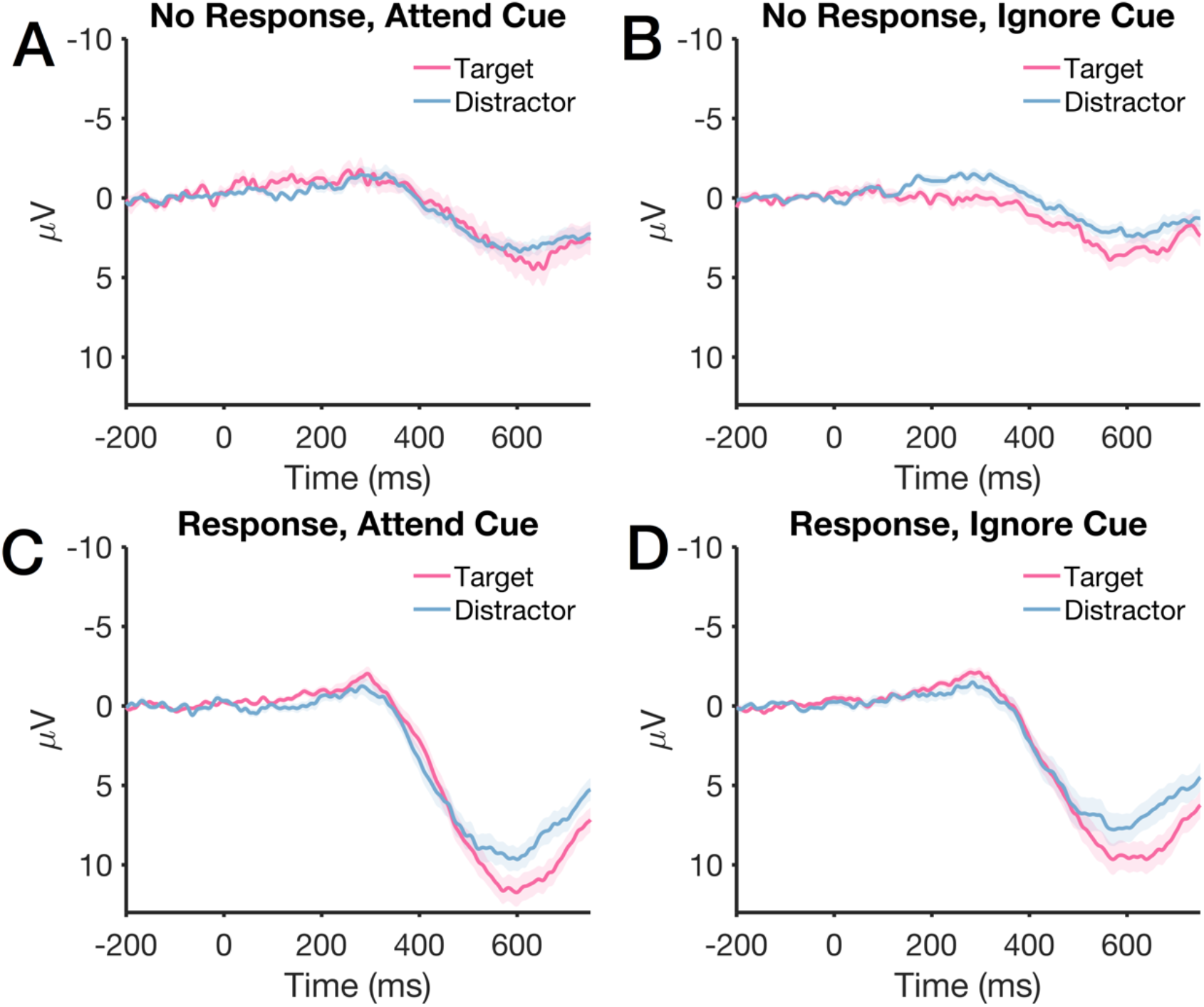
P3 component at electrodes Pz and POz, split by whether or not a response was made. (A) No response made, “attend cue” condition. (B) No response made, “ignore cue” condition. (C) Response made, “attend cue” condition. (D) Response made, “ignore cue” condition. Shaded error bars represent standard error of the mean.

**Appendix P.** Study overview for studies employing a variant of the “competing gratings” task. From left to right, columns indicate: “Exp.” = Experiment number out of those reviewed, “Ref” = Paper reference, “N” = number of subjects in the experiment, “Total Trials” = total number of trials completed by the participant, “Trials Per Cond.” = The number of trials that could be analyzed per condition (i.e., after excluding target and distractor onsets), “Stimulus Duration” = the duration, in seconds, that participants attended the stimulus, “Freq.” = Frequency, in Hertz (Hz), that the stimuli flickered at, “Sig” = Qualitative code for the overall presence of a basic attention effects (when expected); 1 = attended > ignored, −1 = attended < ignored, 0.5 = mixed effects across conditions, 0 = null, n/a = statistical values for the basic attention effect not reported directly. Notes: * Statistics were performed for individuals but not across subjects; standard attention effect in one condition, reversed effect in the other. **Group level statistics not reported. ***No attention effect for the main flicker frequencies (14.2, 17 Hz), but attention effect for the slow oscillating changes to the Gabor’s features (3.14, 3.62 Hz).

| Exp. | Ref. | N | Total Trials | Trials Per Cond. | Stim. Dur. (s) | Freq. (Hz) | Sig. |
| --- | --- | --- | --- | --- | --- | --- | --- |
| 1 | (Chen et al., 2003) | 11 | 16 | 8 | 100 | 7.41, 8.33 | .5* |
| 2 | (Wang et al., 2007),<br>Exp 1 | 12 | 16 | 8 | 120 | 7.14, 8.33 | 0 |
| 3 | (Wang et al., 2007),<br>Exp 2 | 12 | 16 | 8 | 120 | 7.69, 7.14, 8.33 | -1 |
| 4 | (Wang et al., 2007),<br>Exp 3 | 12 | 16 | 8 | 120 | 6.67, 7.14, 7.69, 8.33 | -1 |
| 5 | (Allison et al., 2008) | 14 | 8 | 4 | 60 | 10, 12 | n/a** |
| 6 | (Keitel & Müller, 2016) | 13 | 600 | 75 | 3.5 | 3.14, 3.62, 14.2, 17 | .5*** |

**Appendix Q.** Study overview for studies employing a variant of the “whole-field flicker” task. From left to right, columns indicate: “Exp.” = Experiment number out of those reviewed, “Ref” = Paper reference, “N” = number of subjects in the experiment, “Total Trials” = total number of trials completed by the participant, “Trials Per Cond.” = The number of trials that could be analyzed per condition (i.e., after excluding target and distractor onsets), “Stimulus Duration” = the duration, in seconds, that participants attended the stimulus, “Freq.” = Frequency, in Hertz (Hz), of the stimulus flicker, “Sig” = Qualitative code for the overall presence of a basic attention effects (when expected); 1 = attended > ignored, −1 = attended < ignored, 0.5 = mixed effects across conditions, 0 = null, n/a = statistical values for the basic attention effect not reported directly. Notes: *Analyzed harmonics (2F, 4F) but not the fundamental frequency. 2F but not 4F had a significant attention effect. ** Attention modulation scores were only compared across conditions, not to baseline; they are presumably overall significant, but this was not formally tested.

| Exp. | Ref. | N | Total Trials | Trials Per Cond. | Stim. Dur. (s) | Freq. (Hz) | Sig. |
| --- | --- | --- | --- | --- | --- | --- | --- |
| 7 | (Pei et al., 2002) | 11 | 20 | 20 | 8 | 2.4, 3 | n/a* |
| 8 | (Müller et al., 2006) | 11 | 450 | 153 | 4.114 | 7, 11.67 | 1 |
| 9 | (Andersen et al., 2008) | 15 | 600 | 90 | 3.092 | 10, 12, 15, 17.14 | 1 |
| 10 | (Andersen et al., 2009) | 15 | 432 | 72 | 3.042 | 10, 12 | 1 |
| 11 | (Andersen & Müller, 2010) | 16 | 480 | 240 | 2 | 11.98, 16.77 | 1 |
| 12 | (Quigley et al., 2010) | 10 | 440 | 110 | 2.2 | 8, 12 | 1 |
| 13 | (Zhang et al., 2010) | 18 | 300 | 300 | 4 | 10, 12 | 1 |
| 14 | (Andersen et al., 2012) | 16 | 300 | 60 | 8.5 | 10, 12 | 1 |
| 15 | (Quigley & Müller, 2014) | 20 | 320 | 90 | 4.167 | 15, 17 | 1 |
| 16 | (Andersen et al., 2015) | 15 | 192 | 96 | 15 | 8, 10, 12, 15 | 1 |
| 17 | (Forschack et al., 2017) | 23 | 480 | 120 | 1.783 | 10, 12.5, 15, 17.5 | 1 |
| 18 | (Martinovic & Andersen, 2018) | 9 | 768 | 23 | 6.5 | 10, 12 | n/a** |
| 19 | (Martinovic et al., 2018)<br>Exp 1 | 11 | 600 | 70 | 3.14 | 8.57, 10, 12, 15 | 1 |
| 20 | (Martinovic et al., 2018)<br>Exp 2 | 14 | 600 | 70 | 3.14 | 8.57, 10, 12, 15 | 1 |
| 21 | (Steinhauser & Andersen, 2019) | 17 | 1600 | 400 | 1 | 10, 15 | 1 |

**Appendix R.** Study overview for studies employing a variant of the “hemifield flicker” task. From left to right, columns indicate: “Exp.” = Experiment number out of those reviewed, “Ref” = Paper reference, “N” = number of subjects in the experiment, “Total Trials” = total number of trials completed by the participant, “Trials Per Cond.” = The number of trials that could be analyzed per condition (i.e., after excluding target and distractor onsets), “Stimulus Duration” = the duration, in seconds, that participants attended the stimulus, “Freq.” = Frequency, in Hertz (Hz), of the stimulus flicker, “Sig” = Qualitative code for the overall presence of a basic attention effects (when expected); 1 = attended > ignored, −1 = attended < ignored, 0.5 = mixed effects across conditions, 0 = null, n/a = statistical values for the basic attention effect not reported directly.

| Exp. | Ref. | N | Total Trials | Trials Per Cond. | Stim. Dur. (s) | Freq. (Hz) | Sig. |
| --- | --- | --- | --- | --- | --- | --- | --- |
| 22 | (Andersen et al., 2011) | 19 | 600 | 100 | 3.05 | 8.46, 11.85, 14.81, 19.75 | 1 |
| 23 | (Andersen et al., 2013)<br>Exp 1 | 13 | 560 | 160 | 2.94 | 7.5, 8.57, 10, 12 | 1 |
| 24 | (Andersen et al., 2013)<br>Exp 2 | 11 | 560 | 320 | 2.94 | 7, 8.57, 10, 12 | 1 |
| 25 | (Störmer & Alvarez, 2014) | 16 | 640 | 160 | 2.6 | 7.1, 8.5, 10.7 | 1 |
| 26 | (Müller et al., 2018) | 23 | 480 | 120 | 1.783 | 6.5, 8.5, 11.5, 13.5 | 1 |
| 27 | (Adamian et al., 2019) | 16 | 672 | 128 | 2.94 | 7.5, 8.57, 10, 12 | 1 |

**Appendix S.** Study overview for studies employing a variant of the “attend central, peripheral flicker” task. From left to right, columns indicate: “Exp.” = Experiment number out of those reviewed, “Ref” = Paper reference, “N” = number of subjects in the experiment, “Total Trials” = total number of trials completed by the participant, “Trials Per Cond.” = The number of trials that could be analyzed per condition (i.e., after excluding target and distractor onsets), “Stimulus Duration” = the duration, in seconds, that participants attended the stimulus, “Freq.” = Frequency, in Hertz (Hz), of the stimulus flicker, “Sig” = Qualitative code for the overall presence of a basic attention effects (when expected); 1 = attended > ignored, −1 = attended < ignored, 0.5 = mixed effects across conditions, 0 = null, n/a = statistical values for the basic attention effect not reported directly. Notes: *No attention effect at *a priori* electrode; other electrodes were examined post-hoc, but statistics were not reported for each. **Significant in 1 of 2 expected conditions.

| Exp. | Ref. | N | Total Trials | Trials Per Cond. | Stim. Dur. (s) | Freq. (Hz) | Sig. |
| --- | --- | --- | --- | --- | --- | --- | --- |
| 29 | (Painter et al., 2014)<br>Exp 1 | 20 | 288 | 144 | 7.2 | 12.5, 16.7 | 1 |
| 30 | (Painter et al., 2014)<br>Exp 2 | 20 | 216 | 216 | 8.4 | 7.6, 13.3, 17.8 | 1 |
| 31 | (Painter et al., 2015, p. 2) | 20 | 512 | 128 | 8 | 8, 12 | 0* |
| 32 | (Jiang et al., 2017) | 23 | 288 | 144 | 8.4 | 12, 15 | .5** |
| 33 | (Chu & D’Zmura, 2019)<br>Exp 1 | 20 | 128 | 32 | 7 | 12.5, 18.75 | 1 |
| 34 | (Chu & D’Zmura, 2019)<br>Exp 2 | 21 | 128 | 32 | 9 | 12.5, 18.75 | 1 |

**Appendix T.** Accuracy and task variant for studies where participants detected a target within the flickering stimulus (whole-field and hemifield flicker tasks). To test if the difficulty of our task may have contributed to our null results, we examined behavior from studies in which participants monitored for a target in the flickering stimulus (i.e., whole-field and hemifield flicker tasks). We also noted the type of target and how long it was on the screen. Notes: †Values were not listed in the text, so some values were approximated based on the figures (e.g., hit rates or d’ depicted in a bar graph). *Analyzed harmonics (2F, 4F) but not the fundamental frequency. 2F but not 4F had a significant attention effect. **Attention modulation scores were only compared across conditions, not to baseline; they were presumably significant overall, but this was not formally tested.

| Exp. | Ref. | Behavior | Target type | Dur. (ms) | Sig. |
| --- | --- | --- | --- | --- | --- |
| 7 | (Pei et al., 2002) | n/a | No Targets | n/a | n/a* |
| 8 | (Müller et al., 2006) | $d' = 1.95 - 2.89$ | 75% Coherent Motion | 586 | 1 |
| 9 | (Andersen et al., 2008) | $d' = 2.74 - 3.25$ | 70% Coherent Motion | 500 | 1 |
| 10 | (Andersen et al., 2009) | $d' = 2.67 - 3.23†$ | 20% Luminance Decrement | 200 | 1 |
| 11 | (Andersen & Müller, 2010) | $d' = 1.83$ | 75% Coherent Motion | 298 | 1 |
| 12 | (Quigley et al., 2010) | $d' = 2.665$ | 85% Coherent Motion | 556 | 1 |
| 13 | (Zhang et al., 2010) | n/a | No Targets | n/a | 1 |
| 14 | (Andersen et al., 2012) | $d' = 2.64$ | 50% Coherent Motion | 400 | 1 |
| 15 | (Quigley & Müller, 2014) | Acc = 87.5% - 98%† | 40% Coherent Oblique Motion | 500 | 1 |
| 16 | (Andersen et al., 2015) | $d' = 1.3 - 1.75†$ | 70% Coherent Motion | 500 | 1 |
| 17 | (Forschack et al., 2017) | $d' = 2$ | 60% Coherent Motion | 300 | 1 |
| 18 | (Martinovic & Andersen, 2018) | $d' = 0.8 - 3.0†$ | 50% Coherent Motion | 400 | n/a** |
| 19 | (Martinovic et al., 2018)<br>Exp 1 | $d' = 1.05$ | 50% Coherent Motion | 400 | 1 |
| 20 | (Martinovic et al., 2018)<br>Exp 2 | $d' = 1.0$ | 50% Coherent Motion | 400 | 1 |
| 21 | (Steinhauser & Andersen, 2019) | Acc = 90.3% | 75% Coherent Motion | 500 | 1 |
| 22 | (Andersen et al., 2011) | $d' = 0.95 - 2.8$ | 75% Coherent Motion | 500 | 1 |
| 23 | (Andersen et al., 2013)<br>Exp 1 | $d' = 2.133 - 3.111$ | 20% Luminance Decrement | 200 | 1 |
| 24 | (Andersen et al., 2013)<br>Exp 2 | $d' = 2.637$ | 20% Luminance Decrement | 200 | 1 |
| 25 | (Störmer & Alvarez, 2014) | Acc = 78% | 80% Coherent Motion | 230 | 1 |
| 26 | (Müller et al., 2018) | $d' = 1.81$ | 60% Coherent Motion | 300 | 1 |
| 27 | (Adamian et al., 2019) | $d' = 2.8†$ | 20% Luminance Decrement | 200 | 1 |

## Footnotes

* As we did not pre-register Bayesian analysis choices (e.g., choices about priors, etc.), we did not calculate a Bayes Factor for this pre-registered analysis. However, the post-hoc power analysis gives a sense of the degree to which this is a null effect.

## References

Adam, K. C. S., & Serences, J. T. (2020). History-driven modulations of population codes in early visual cortex during visual search [Preprint]. bioRxiv. https://doi.org/10.1101/2020.09.30.321729

Adamian, N., Slaustaite, E., & Andersen, S. K. (2019). Top–Down Attention Is Limited Within but Not Between Feature Dimensions. Journal of Cognitive Neuroscience, 31(8), 1173–1183. https://doi.org/10.1162/jocn_a_01383

Adrian, E. D., & Matthews, B. H. C. (1934). The Berger Rhythm: Potential Changes from the Occiptial Lobes in Man. Brain, 57(4), 355–385.

Allison, B. Z., McFarland, D. J., Schalk, G., Zheng, S. D., Jackson, M. M., & Wolpaw, J. R. (2008). Towards an independent brain–computer interface using steady state visual evoked potentials. Clinical Neurophysiology, 119(2), 399–408. https://doi.org/10.1016/j.clinph.2007.09.121

Andersen, S. K., Fuchs, S., & Müller, M. M. (2011). Effects of Feature-selective and Spatial Attention at Different Stages of Visual Processing. Journal of Cognitive Neuroscience, 23(1), 238–246. https://doi.org/10.1162/jocn.2009.21328

Andersen, S. K., Hillyard, S. A., & Müller, M. M. (2008). Attention Facilitates Multiple Stimulus Features in Parallel in Human Visual Cortex. Current Biology, 18(13), 1006–1009. https://doi.org/10.1016/j.cub.2008.06.030

Andersen, S. K., Hillyard, S. A., & Müller, M. M. (2013). Global Facilitation of Attended Features Is Obligatory and Restricts Divided Attention. Journal of Neuroscience, 33(46), 18200–18207. https://doi.org/10.1523/JNEUROSCI.1913-13.2013

Andersen, S. K., & Müller, M. M. (2010). Behavioral performance follows the time course of neural facilitation and suppression during cued shifts of feature-selective attention. Proceedings of the National Academy of Sciences, 107(31), 13878–13882. https://doi.org/10.1073/pnas.1002436107

Andersen, S. K., Müller, M. M., & Hillyard, S. A. (2009). Color-selective attention need not be mediated by spatial attention. Journal of Vision, 9(6), 2–2. https://doi.org/10.1167/9.6.2

Andersen, S. K., Müller, M. M., & Hillyard, S. A. (2011). Tracking the allocation of attention in visual scenes with steady-state evoked potentials. In M. I. Posner (Ed.), Cognitive Neuroscience of Attention (2nd ed.). The Guilford Press.

Andersen, S. K., Müller, M. M., & Hillyard, S. A. (2015). Attentional Selection of Feature Conjunctions Is Accomplished by Parallel and Independent Selection of Single Features. Journal of Neuroscience, 35(27), 9912–9919. https://doi.org/10.1523/JNEUROSCI.5268-14.2015

Andersen, S. K., Müller, M. M., & Martinovic, J. (2012). Bottom-Up Biases in Feature-Selective Attention. Journal of Neuroscience, 32(47), 16953–16958. https://doi.org/10.1523/JNEUROSCI.1767-12.2012

Antonov, P. A., Chakravarthi, R., & Andersen, S. K. (2020). Too little, too late, and in the wrong place: Alpha band activity does not reflect an active mechanism of selective attention. NeuroImage, 219, 117006. https://doi.org/10.1016/j.neuroimage.2020.117006

Appelbaum, L. G., & Norcia, A. M. (2009). Attentive and pre-attentive aspects of figural processing. Journal of Vision, 9(11), 18–18. https://doi.org/10.1167/9.11.18

Arita, J. T., Carlisle, N. B., & Woodman, G. F. (2012). Templates for rejection: Configuring attention to ignore task-irrelevant features. Journal of Experimental Psychology: Human Perception and Performance, 38(3), 580–584. https://doi.org/10.1037/a0027885

Awh, E., Belopolsky, A. V., & Theeuwes, J. (2012). Top-down versus bottom-up attentional control: A failed theoretical dichotomy. Trends in Cognitive Sciences, 16(8), 437–443. https://doi.org/10.1016/j.tics.2012.06.010

Baker, D. H., Vilidaite, G., Lygo, F. A., Smith, A. K., Flack, T. R., Gouws, A. D., & Andrews, T. J. (2019). Power contours: Optimising sample size and precision in experimental psychology and human neuroscience. ArXiv:1902.06122[q-Bio, Stat]. http://arxiv.org/abs/1902.06122

Bartsch, M. V., Loewe, K., Merkel, C., Heinze, H.-J., Schoenfeld, M. A., Tsotsos, J. K., & Hopf, J.-M. (2017). Attention to Color Sharpens Neural Population Tuning via Feedback Processing in the Human Visual Cortex Hierarchy. The Journal of Neuroscience, 37(43), 10346–10357. https://doi.org/10.1523/JNEUROSCI.0666-17.2017

Beck, V. M., & Hollingworth, A. (2015). Evidence for negative feature guidance in visual search is explained by spatial recoding. Journal of Experimental Psychology: Human Perception and Performance, 41(5), 1190–1196. https://doi.org/10.1037/xhp0000109

Becker, M. W., Hemsteger, S., & Peltier, C. (2015). No templates for rejection: A failure to configure attention to ignore task-irrelevant features. Visual Cognition, 23(9–10), 1150–1167. https://doi.org/10.1080/13506285.2016.1149532

Boudewyn, M. A., Luck, S. J., Farrens, J. L., & Kappenman, E. S. (2018). How many trials does it take to get a significant ERP effect? It depends. Psychophysiology, 55(6), e13049. https://doi.org/10.1111/psyp.13049

Boylan, M. R., Kelly, M. N., Thigpen, N. N., & Keil, A. (2019). Attention to a threat-related feature does not interfere with concurrent attentive feature selection. Psychophysiology, 56(6), e13332. https://doi.org/10.1111/psyp.13332

Brainard, D. H. (1997). The Psychophysics Toolbox. Spatial Vision, 10(4), 433–436. https://doi.org/10.1163/156856897X00357

Bridwell, D. A., & Srinivasan, R. (2012). Distinct Attention Networks for Feature Enhancement and Suppression in Vision. Psychological Science, 23(10), 1151–1158. https://doi.org/10.1177/0956797612440099

Button, K. S., Ioannidis, J. P. A., Mokrysz, C., Nosek, B. A., Flint, J., Robinson, E. S. J., & Munafò, M. R. (2013). Power failure: Why small sample size undermines the reliability of neuroscience. Nature Reviews Neuroscience, 14(5), 365–376. https://doi.org/10.1038/nrn3475

Button, K. S., & Munafò, M. R. (2017). Powering Reproducible Research. In S. O. Lilienfeld & I. D. Waldman (Eds.), Psychological Science Under Scrutiny (pp. 22–33). John Wiley & Sons, Inc. https://doi.org/10.1002/9781119095910.ch2

Canolty, R. T., Edwards, E., Dalal, S. S., Soltani, M., Nagarajan, S. S., Kirsch, H. E., Berger, M. S., Barbaro, N. M., & Knight, R. T. (2006). High Gamma Power Is Phase-Locked to Theta Oscillations in Human Neocortex. Science, 313(5793), 1626–1628. https://doi.org/10.1126/science.1128115

Carlisle, N. B., & Nitka, A. W. (2019). Location-based explanations do not account for active attentional suppression. Visual Cognition, 27(3–4), 305–316. https://doi.org/10.1080/13506285.2018.1553222

Chang, S., & Egeth, H. E. (2019). Enhancement and Suppression Flexibly Guide Attention. Psychological Science, 30(12), 1724–1732. https://doi.org/10.1177/0956797619878813

Chapman, A. F., Geweke, F., & Störmer, V. S. (2019). Feature-based attention resolves differences in target-distractor similarity through multiple mechanisms. Journal of Vision, 19(10), 45a. https://doi.org/10.1167/19.10.45a

Chen, Y., Seth, A. K., Gally, J. A., & Edelman, G. M. (2003). The power of human brain magnetoencephalographic signals can be modulated up or down by changes in an attentive visual task. Proceedings of the National Academy of Sciences, 100(6), 3501–3506. https://doi.org/10.1073/pnas.0337630100

Chu, V. C., & D’Zmura, M. (2019). Tracking feature-based attention. Journal of Neural Engineering, 16(1), 016022. https://doi.org/10.1088/1741-2552/aaed17

Clayson, P. E., Carbine, K. A., Baldwin, S. A., & Larson, M. J. (2019). Methodological reporting behavior, sample sizes, and statistical power in studies of event-related potentials: Barriers to reproducibility and replicability. Psychophysiology, 56(11). https://doi.org/10.1111/psyp.13437

Clementz, B. A., Wang, J., & Keil, A. (2008). Normal Electrocortical Facilitation But Abnormal Target Identification during Visual Sustained Attention in Schizophrenia. Journal of Neuroscience, 28(50), 13411–13418. https://doi.org/10.1523/JNEUROSCI.4095-08.2008

Cohen, M. R., & Maunsell, J. H. R. (2011). Using Neuronal Populations to Study the Mechanisms Underlying Spatial and Feature Attention. Neuron, 70(6), 1192–1204. https://doi.org/10.1016/j.neuron.2011.04.029

Cohen, M. X., & Gulbinaite, R. (2017). Rhythmic entrainment source separation: Optimizing analyses of neural responses to rhythmic sensory stimulation. NeuroImage, 147, 43–56. https://doi.org/10.1016/j.neuroimage.2016.11.036

Combrisson, E., & Jerbi, K. (2015). Exceeding chance level by chance: The caveat of theoretical chance levels in brain signal classification and statistical assessment of decoding accuracy. Journal of Neuroscience Methods, 250, 126–136. https://doi.org/10.1016/j.jneumeth.2015.01.010

Conci, M., Deichsel, C., Müller, H. J., & Töllner, T. (2019). Feature guidance by negative attentional templates depends on search difficulty. Visual Cognition, 27(3–4), 317–326. https://doi.org/10.1080/13506285.2019.1581316

Cooper, H., DeNeve, K., & Charlton, K. (1997). Finding the missing science: The fate of studies submitted for review by a human subjects committee. Psychological Methods, 2(4), 447–452. https://doi.org/10.1037/1082-989X.2.4.447

Cunningham, C. A., & Egeth, H. E. (2016). Taming the White Bear: Initial Costs and Eventual Benefits of Distractor Inhibition. Psychological Science, 27(4), 476–485. https://doi.org/10.1177/0956797615626564

Dickersin, Kay. (1990). The Existence of Publication Bias and Risk Factors for Its Occurrence. JAMA: The Journal of the American Medical Association, 263(10), 1385. https://doi.org/10.1001/jama.1990.03440100097014

Dickersin, Kay, Min, Y. I., & Meinert, C. L. (1992). Factors influencing publication of research results. Follow-up of applications submitted to two institutional review boards. JAMA, 267(3), 374–378.

Ding, J., Sperling, G., & Srinivasan, R. (2006). Attentional Modulation of SSVEP Power Depends on the Network Tagged by the Flicker Frequency. Cerebral Cortex, 16(7), 1016–1029. https://doi.org/10.1093/cercor/bhj044

Donohue, S. E., Bartsch, M. V., Heinze, H.-J., Schoenfeld, M. A., & Hopf, J.-M. (2018). Cortical Mechanisms of Prioritizing Selection for Rejection in Visual Search. The Journal of Neuroscience, 38(20), 4738–4748. https://doi.org/10.1523/JNEUROSCI.2407-17.2018

Dwan, K., Altman, D. G., Arnaiz, J. A., Bloom, J., Chan, A.-W., Cronin, E., Decullier, E., Easterbrook, P. J., Von Elm, E., Gamble, C., Ghersi, D., Ioannidis, J. P. A., Simes, J., & Williamson, P. R. (2008). Systematic Review of the Empirical Evidence of Study Publication Bias and Outcome Reporting Bias. PLoS ONE, 3(8), e3081. https://doi.org/10.1371/journal.pone.0003081

Failing, M., Feldmann-Wüstefeld, T., Wang, B., Olivers, C., & Theeuwes, J. (2019). Statistical regularities induce spatial as well as feature-specific suppression. Journal of Experimental Psychology: Human Perception and Performance, 45(10), 1291–1303. https://doi.org/10.1037/xhp0000660

Ferguson, C. J., & Heene, M. (2012). A Vast Graveyard of Undead Theories: Publication Bias and Psychological Science’s Aversion to the Null. Perspectives on Psychological Science, 7(6), 555–561. https://doi.org/10.1177/1745691612459059

Forschack, N., Andersen, S. K., & Müller, M. M. (2017). Global Enhancement but Local Suppression in Feature-based Attention. Journal of Cognitive Neuroscience, 29(4), 619–627. https://doi.org/10.1162/jocn_a_01075

Franco, A., Malhotra, N., & Simonovits, G. (2014). Social science. Publication bias in the social sciences: Unlocking the file drawer. Science (New York, N.Y.), 345(6203), 1502–1505. https://doi.org/10.1126/science.1255484

Garcia, J. O., Srinivasan, R., & Serences, J. T. (2013). Near-Real-Time Feature-Selective Modulations in Human Cortex. Current Biology, 23(6), 515–522. https://doi.org/10.1016/j.cub.2013.02.013

Geng, J. J. (2014). Attentional Mechanisms of Distractor Suppression. Current Directions in Psychological Science, 23(2), 147–153. https://doi.org/10.1177/0963721414525780

Geng, J. J., Won, B.-Y., & Carlisle, N. B. (2019). Distractor Ignoring: Strategies, Learning, and Passive Filtering. Current Directions in Psychological Science, 28(6), 600–606. https://doi.org/10.1177/0963721419867099

Geweke, F., Li, S.-C., & Störmer, V. (2018). Feature-based attention is constrained to attended locations in older adults. Journal of Vision, 18(10), 306. https://doi.org/10.1167/18.10.306

Guan, M., & Vandekerckhove, J. (2016). A Bayesian approach to mitigation of publication bias. Psychonomic Bulletin & Review, 23(1), 74–86. https://doi.org/10.3758/s13423-015-0868-6

Gulbinaite, R., Roozendaal, D. H. M., & VanRullen, R. (2019). Attention differentially modulates the amplitude of resonance frequencies in the visual cortex. NeuroImage, 203, 116146. https://doi.org/10.1016/j.neuroimage.2019.116146

Hasan, R., Srinivasan, R., & Grossman, E. D. (2017). Feature-based attentional tuning during biological motion detection measured with SSVEP. Journal of Vision, 17(9), 22. https://doi.org/10.1167/17.9.22

Haxby, J., Horwitz, B., Ungerleider, L., Maisog, J., Pietrini, P., & Grady, C. (1994). The functional organization of human extrastriate cortex: A PET-rCBF study of selective attention to faces and locations. The Journal of Neuroscience, 14(11), 6336–6353. https://doi.org/10.1523/JNEUROSCI.14-11-06336.1994

Herrmann, C. S. (2001). Human EEG responses to 1?100□Hz flicker: Resonance phenomena in visual cortex and their potential correlation to cognitive phenomena. Experimental Brain Research, 137(3-4), 346–353. https://doi.org/10.1007/s002210100682

Ipata, A. E., Gee, A. L., Gottlieb, J., Bisley, J. W., & Goldberg, M. E. (2006). LIP responses to a popout stimulus are reduced if it is overtly ignored. Nature Neuroscience, 9(8), 1071–1076. https://doi.org/10.1038/nn1734

Itthipuripat, S., Deering, S., & Serences, J. T. (2019). When Conflict Cannot be Avoided: Relative Contributions of Early Selection and Frontal Executive Control in Mitigating Stroop Conflict. Cerebral Cortex, 29(12), 5037–5048. https://doi.org/10.1093/cercor/bhz042

Itthipuripat, S., Garcia, J. O., & Serences, J. T. (2013). Temporal dynamics of divided spatial attention. Journal of Neurophysiology, 109(9), 2364–2373. https://doi.org/10.1152/jn.01051.2012

Jiang, Y., Wu, X., & Gao, X. (2017). A category-specific top-down attentional set can affect the neural responses outside the current focus of attention. Neuroscience Letters, 659, 80–85. https://doi.org/10.1016/j.neulet.2017.07.029

Kadel, H., Feldmann-Wüstefeld, T., & Schubö, A. (2017). Selection history alters attentional filter settings persistently and beyond top-down control: Selection history alters attentional filter settings. Psychophysiology, 54(5), 736–754. https://doi.org/10.1111/psyp.12830

Kaiser, P. K. (1988). Sensation luminance: A new name to distinguish CIE luminance from luminance dependent on an individual’s spectral sensitivity. Vision Research, 28(3), 455–456. https://doi.org/10.1016/0042-6989(88)90186-1

Kappenman, E. S., & Luck, S. J. (2010). The effects of electrode impedance on data quality and statistical significance in ERP recordings. Psychophysiology. https://doi.org/10.1111/j.1469-8986.2010.01009.x

Kashiwase, Y., Matsumiya, K., Kuriki, I., & Shioiri, S. (2012). Time Courses of Attentional Modulation in Neural Amplification and Synchronization Measured with Steady-state Visual-evoked Potentials. Journal of Cognitive Neuroscience, 24(8), 1779–1793. https://doi.org/10.1162/jocn_a_00212

Kastner, S., & Ungerleider, L. G. (2000). Mechanisms of Visual Attention in the Human Cortex. Annual Review of Neuroscience, 23(1), 315–341. https://doi.org/10.1146/annurev.neuro.23.1.315

Keitel, C., & Müller, M. M. (2016). Audio-visual synchrony and feature-selective attention co-amplify early visual processing. Experimental Brain Research, 234(5), 1221–1231. https://doi.org/10.1007/s00221-015-4392-8

Kim, Y. J., Grabowecky, M., Paller, K. A., Muthu, K., & Suzuki, S. (2007). Attention induces synchronization-based response gain in steady-state visual evoked potentials. Nature Neuroscience, 10(1), 117–125. https://doi.org/10.1038/nn1821

Kiyonaga, A., & Egner, T. (2016). Center-Surround Inhibition in Working Memory. Current Biology, 26(1), 64–68. https://doi.org/10.1016/j.cub.2015.11.013

Kleiner, M., Brainard, D., & Pelli, D. (2007). What’s new in Psychtoolbox-3? European Conference on Visual Perception (ECVP), Arezzo, Italy. https://pdfs.semanticscholar.org/04d4/7572cec08b7a582a9366e5ac61dcfd633f2a.pdf

Kravitz, D., & Mitroff, S. (2017). Estimates of a priori power and false discovery rates induced by post-hoc changes from thousands of independent replications. Journal of Vision, 17(10), 223. https://doi.org/10.1167/17.10.223

Lamy, D. F., Antebi, C., Aviani, N., & Carmel, T. (2008). Priming of Pop-out provides reliable measures of target activation and distractor inhibition in selective attention. Vision Research, 48(1), 30–41. https://doi.org/10.1016/j.visres.2007.10.009

Lamy, D. F., & Kristjansson, A. (2013). Is goal-directed attentional guidance just intertrial priming? A review. Journal of Vision, 13(3), 14–14. https://doi.org/10.1167/13.3.14

Lithari, C., Sánchez-García, C., Ruhnau, P., & Weisz, N. (2016). Large-scale network-level processes during entrainment. Brain Research, 1635, 143–152. https://doi.org/10.1016/j.brainres.2016.01.043

Luck, S. J. (2005). An Introduction to the Event-Related Potential Technique (1st ed.). MIT Press.

Martinez-Trujillo, J. C., & Treue, S. (2004). Feature-Based Attention Increases the Selectivity of Population Responses in Primate Visual Cortex. Current Biology, 14(9), 744–751. https://doi.org/10.1016/j.cub.2004.04.028

Martinovic, J., & Andersen, S. K. (2018). Cortical summation and attentional modulation of combined chromatic and luminance signals. NeuroImage, 176, 390–403. https://doi.org/10.1016/j.neuroimage.2018.04.066

Martinovic, J., Wuerger, S. M., Hillyard, S. A., Müller, M. M., & Andersen, S. K. (2018). Neural mechanisms of divided feature-selective attention to colour. NeuroImage, 181, 670–682. https://doi.org/10.1016/j.neuroimage.2018.07.033

Mishkin, M., & Ungerleider, L. G. (1982). Contribution of striate inputs to the visuospatial functions of parieto-preoccipital cortex in monkeys. Behavioural Brain Research, 6(1), 57–77. https://doi.org/10.1016/0166-4328(82)90081-X

Moher, J., & Egeth, H. E. (2012). The ignoring paradox: Cueing distractor features leads first to selection, then to inhibition of to-be-ignored items. Attention, Perception, & Psychophysics, 74(8), 1590–1605. https://doi.org/10.3758/s13414-012-0358-0

Moher, J., Lakshmanan, B. M., Egeth, H. E., & Ewen, J. B. (2014). Inhibition Drives Early Feature-Based Attention. Psychological Science, 25(2), 315–324. https://doi.org/10.1177/0956797613511257

Morgan, S. T., Hansen, J. C., & Hillyard, S. A. (1996). Selective attention to stimulus location modulates the steady-state visual evoked potential. Proceedings of the National Academy of Sciences of the United States of America, 93(10), 4770–4774.

Müller, M. M., Andersen, S., Trujillo, N. J., Valdes-Sosa, P., Malinowski, P., & Hillyard, S. A. (2006). Feature-selective attention enhances color signals in early visual areas of the human brain. Proceedings of the National Academy of Sciences, 103(38), 14250–14254. https://doi.org/10.1073/pnas.0606668103

Müller, M. M., Gundlach, C., Forschack, N., & Brummerloh, B. (2018). It takes two to tango: Suppression of task-irrelevant features requires (spatial) competition. NeuroImage, 178, 485–492. https://doi.org/10.1016/j.neuroimage.2018.05.073

Müller, M. M., & Hillyard, S. (2000). Concurrent recording of steady-state and transient event-related potentials as indices of visual-spatial selective attention. Clinical Neurophysiology, 111(9), 1544–1552. https://doi.org/10.1016/S1388-2457(00)00371-0

Müller, M. M., Picton, T. W., Valdes-Sosa, P., Riera, J., Teder-Sälejärvi, W. A., & Hillyard, S. A. (1998). Effects of spatial selective attention on the steady-state visual evoked potential in the 20-28 Hz range. Cognitive Brain Research, 6(4), 249–261. https://doi.org/10.1016/S0926-6410(97)00036-0

Müller, M. M., Teder-Sälejärvi, W., & Hillyard, S. A. (1998). The time course of cortical facilitation during cued shifts of spatial attention. Nature Neuroscience, 1(7), 631–634. https://doi.org/10.1038/2865

Norcia, A. M., Appelbaum, L. G., Ales, J. M., Cottereau, B. R., & Rossion, B. (2015). The steady-state visual evoked potential in vision research: A review. Journal of Vision, 15(6), 4. https://doi.org/10.1167/15.6.4

Nunez, M. D., Srinivasan, R., & Vandekerckhove, J. (2015). Individual differences in attention influence perceptual decision making. Frontiers in Psychology, 8, 18. https://doi.org/10.3389/fpsyg.2015.00018

Owen, A. M., Milner, B., Petrides, M., & Evans, A. C. (1996). Memory for object features versus memory for object location: A positron-emission tomography study of encoding and retrieval processes. Proceedings of the National Academy of Sciences, 93(17), 9212–9217. https://doi.org/10.1073/pnas.93.17.9212

Painter, D. R., Dux, P. E., & Mattingley, J. B. (2015). Causal involvement of visual area MT in global feature-based enhancement but not contingent attentional capture. NeuroImage, 118, 90–102. https://doi.org/10.1016/j.neuroimage.2015.06.019

Painter, D. R., Dux, P. E., Travis, S. L., & Mattingley, J. B. (2014). Neural Responses to Target Features outside a Search Array Are Enhanced during Conjunction but Not Unique-Feature Search. Journal of Neuroscience, 34(9), 3390–3401. https://doi.org/10.1523/JNEUROSCI.3630-13.2014

Pei, F., Pettet, M. W., & Norcia, A. M. (2002). Neural correlates of object-based attention. Journal of Vision, 2(9), 1. https://doi.org/10.1167/2.9.1

Pelli, D. G. (1997). The VideoToolbox software for visual psychophysics: Transforming numbers into movies. Spatial Vision, 10(4), 437–442. https://doi.org/10.1163/156856897X00366

Quigley, C., Andersen, S. K., Schulze, L., Grunwald, M., & Müller, M. M. (2010). Feature-selective attention: Evidence for a decline in old age. Neuroscience Letters, 474(1), 5–8. https://doi.org/10.1016/j.neulet.2010.02.053

Quigley, C., & Müller, M. M. (2014). Feature-Selective Attention in Healthy Old Age: A Selective Decline in Selective Attention? Journal of Neuroscience, 34(7), 2471–2476. https://doi.org/10.1523/JNEUROSCI.2718-13.2014

Reeder, R. R., Olivers, C. N. L., & Pollmann, S. (2017). Cortical evidence for negative search templates. Visual Cognition, 25(1–3), 278–290. https://doi.org/10.1080/13506285.2017.1339755

Regan, D. (1977). Steady-state evoked potentials. Journal of the Optical Society of America, 67(11), 1475. https://doi.org/10.1364/JOSA.67.001475

Rosenthal, R. (1979). The file drawer problem and tolerance for null results. Psychological Bulletin, 86(3), 638–641. https://doi.org/10.1037/0033-2909.86.3.638

Rungratsameetaweemana, N., Itthipuripat, S., Salazar, A., & Serences, J. T. (2018). Expectations Do Not Alter Early Sensory Processing during Perceptual Decision-Making. The Journal of Neuroscience, 38(24), 5632–5648. https://doi.org/10.1523/JNEUROSCI.3638-17.2018

Sàenz, M., Buracas, G. T., & Boynton, G. M. (2002). Global effects of feature-based attention in human visual cortex. Nature Neuroscience, 5(7), 631–632. https://doi.org/10.1038/nn876

Sàenz, M., Buraĉas, G. T., & Boynton, G. M. (2003). Global feature-based attention for motion and color. Vision Research, 43(6), 629–637. https://doi.org/10.1016/S0042-6989(02)00595-3

Sawaki, R., & Luck, S. J. (2010). Capture versus suppression of attention by salient singletons: Electrophysiological evidence for an automatic attend-to-me signal. Attention, Perception, & Psychophysics, 72(6), 1455–1470. https://doi.org/10.3758/APP.72.6.1455

Srinivasan, R., Bibi, F. A., & Nunez, P. L. (2006). Steady-State Visual Evoked Potentials: Distributed Local Sources and Wave-Like Dynamics Are Sensitive to Flicker Frequency. Brain Topography, 18(3), 167–187. https://doi.org/10.1007/s10548-006-0267-4

Steinhauser, M., & Andersen, S. K. (2019). Rapid adaptive adjustments of selective attention following errors revealed by the time course of steady-state visual evoked potentials. NeuroImage, 186, 83–92. https://doi.org/10.1016/j.neuroimage.2018.10.059

Sterling, T. D. (1959). Publication Decisions and their Possible Effects on Inferences Drawn from Tests of Significance—Or Vice Versa. Journal of the American Statistical Association, 54(285), 30–34. https://doi.org/10.1080/01621459.1959.10501497

Sterling, T. D., Rosenbaum, W. L., & Weinkam, J. J. (1995). Publication Decisions Revisited: The Effect of the Outcome of Statistical Tests on the Decision to Publish and Vice Versa. The American Statistician, 49(1), 108–112. https://doi.org/10.1080/00031305.1995.10476125

Stilwell, B. T., & Vecera, S. P. (2019). Cued distractor rejection disrupts learned distractor rejection. Visual Cognition, 27(3–4), 327–342. https://doi.org/10.1080/13506285.2018.1564808

Störmer, V. S., & Alvarez, G. A. (2014). Feature-Based Attention Elicits Surround Suppression in Feature Space. Current Biology, 24(17), 1985–1988. https://doi.org/10.1016/j.cub.2014.07.030

Tallon-Baudry, C., Bertrand, O., Delpuech, C., & Pernier, J. (1996). Stimulus Specificity of Phase-Locked and Non-Phase-Locked 40 Hz Visual Responses in Human. The Journal of Neuroscience, 16(13), 4240–4249. https://doi.org/10.1523/JNEUROSCI.16-13-04240.1996

Talsma, D., Doty, T. J., Strowd, R., & Woldorff, M. G. (2006). Attentional capacity for processing concurrent stimuli is larger across sensory modalities than within a modality. Psychophysiology, 43(6), 541–549. https://doi.org/10.1111/j.1469-8986.2006.00452.x

Theeuwes, J. (2013). Feature-based attention: It is all bottom-up priming. Philosophical Transactions of the Royal Society B: Biological Sciences, 368(1628), 20130055. https://doi.org/10.1098/rstb.2013.0055

Thigpen, N. N., Kappenman, E. S., & Keil, A. (2017). Assessing the internal consistency of the event-related potential: An example analysis: Assessing internal consistency of the ERP. Psychophysiology, 54(1), 123–138. https://doi.org/10.1111/psyp.12629

Thigpen, N. N., Petro, N. M., Oschwald, J., Oberauer, K., & Keil, A. (2019). Selection of Visual Objects in Perception and Working Memory One at a Time. Psychological Science, 30(9), 1259–1272. https://doi.org/10.1177/0956797619854067

Toffanin, P., de Jong, R., Johnson, A., & Martens, S. (2009). Using frequency tagging to quantify attentional deployment in a visual divided attention task. International Journal of Psychophysiology, 72(3), 289–298. https://doi.org/10.1016/j.ijpsycho.2009.01.006

Van Moorselaar, D., & Slagter, H. A. (2020). Inhibition in selective attention. Annals of the New York Academy of Sciences, 1464(1), 204–221.

Vatterott, D. B., & Vecera, S. P. (2012). Experience-dependent attentional tuning of distractor rejection. Psychonomic Bulletin & Review, 19(5), 871–878. https://doi.org/10.3758/s13423-012-0280-4

Verghese, P., Kim, Y.-J., & Wade, A. R. (2012). Attention Selects Informative Neural Populations in Human V1. Journal of Neuroscience, 32(46), 16379–16390. https://doi.org/10.1523/JNEUROSCI.1174-12.2012

Vialatte, F.-B., Maurice, M., Dauwels, J., & Cichocki, A. (2010). Steady-state visually evoked potentials: Focus on essential paradigms and future perspectives. Progress in Neurobiology, 90(4), 418–438. https://doi.org/10.1016/j.pneurobio.2009.11.005

Vissers, M. E., Gulbinaite, R., van den Bos, T., & Slagter, H. A. (2017). Protecting visual short-term memory during maintenance: Attentional modulation of target and distractor representations. Scientific Reports, 7(1), 4061. https://doi.org/10.1038/s41598-017-03995-0

Wang, B., & Theeuwes, J. (2018a). Statistical regularities modulate attentional capture. Journal of Experimental Psychology: Human Perception and Performance, 44(1), 13–17. https://doi.org/10.1037/xhp0000472

Wang, B., & Theeuwes, J. (2018b). How to inhibit a distractor location? Statistical learning versus active, top-down suppression. Attention, Perception, & Psychophysics. https://doi.org/10.3758/s13414-018-1493-z

Wang, J., Clementz, B. A., & Keil, A. (2007). The neural correlates of feature-based selective attention when viewing spatially and temporally overlapping images. Neuropsychologia, 45(7), 1393–1399. https://doi.org/10.1016/j.neuropsychologia.2006.10.019

Wang, Y., Miller, J., & Liu, T. (2015). Suppression effects in feature-based attention. Journal of Vision, 15(5), 15. https://doi.org/10.1167/15.5.15

White, A. L., & Carrasco, M. (2011). Feature-based attention involuntarily and simultaneously improves visual performance across locations. Journal of Vision, 11(6), 15–15. https://doi.org/10.1167/11.6.15

Williams, R. S., Pratt, J., & Ferber, S. (2020). Directed avoidance and its effect on visual working memory. Cognition, 201, 104277. https://doi.org/10.1016/j.cognition.2020.104277

Won, B.-Y., & Geng, J. J. (2020). Passive exposure attenuates distraction during visual search. Journal of Experimental Psychology: General. https://doi.org/10.1037/xge0000760

Won, D.-O., Hwang, H.-J., Dähne, S., Müller, K.-R., & Lee, S.-W. (2016). Effect of higher frequency on the classification of steady-state visual evoked potentials. Journal of Neural Engineering, 13(1), 016014. https://doi.org/10.1088/1741-2560/13/1/016014

Zar, J. H. (2010). Biostatistical Analysis (5th ed.). Prentice-Hall.

Zhang, D., Maye, A., Gao, X., Hong, B., Engel, A. K., & Gao, S. (2010). An independent brain–computer interface using covert non-spatial visual selective attention. Journal of Neural Engineering, 7(1), 016010. https://doi.org/10.1088/1741-2560/7/1/016010

Zhang, W., & Luck, S. J. (2009). Feature-based attention modulates feedforward visual processing. Nature Neuroscience, 12(1), 24–25. https://doi.org/10.1038/nn.2223

Zhang, Z., Gaspelin, N., & Carlisle, N. B. (2020). Probing early attention following negative and positive templates. Attention, Perception, & Psychophysics, 82(3), 1166–1175. https://doi.org/10.3758/s13414-019-01864-8

Zhu, D., Bieger, J., Garcia Molina, G., & Aarts, R. M. (2010). A Survey of Stimulation Methods Used in SSVEP-Based BCIs. Computational Intelligence and Neuroscience, 2010, 1–12. https://doi.org/10.1155/2010/702357

